# Collagen has a unique SEC24 preference for efficient export from the endoplasmic reticulum

**DOI:** 10.1101/2021.02.23.431880

**Authors:** Chung-Ling Lu, Steven Ortmeier, Jon Brudvig, Tamara Moretti, Jacob Cain, Simeon A. Boyadjiev, Jill M. Weimer, Jinoh Kim

**Author notes:** To who correspondence should be addressed: Jinoh Kim, 2086 Vet Med, 1800 Christensen Drive, Iowa State University, Ames, IA 50011. **Synopsis:** *SEC24D* gene mutations cause predominantly bone defects. A prevailing explanation is that SEC24D sorts procollagen. Using *Sec24d* KO mice and cultured cells, we show that SEC24A and SEC24B also contribute to ER export of procollagen. In contrast, fibronectin 1 requires either SEC24C or SEC24D for ER export. This finding is significant as it forms the basis for tissue-specific and cargo-specific utilization of SEC24 paralogs for ER export.

## Abstract

SEC24 is mainly involved in cargo sorting during COPII vesicle assembly. There are four SEC24 paralogs (A to D) in mammals, which are classified into two subgroups (SEC24A/B and SEC24C/D). Pathological mutations in *SEC24D* cause osteogenesis imperfecta with craniofacial dysplasia in humans. *sec24d* mutant fish also recapitulate the phenotypes. Consistent with the skeletal phenotypes, the secretion of collagen was severely defective in mutant fish, emphasizing the importance of SEC24D in collagen secretion. However, SEC24D patient-derived fibroblasts show only a mild secretion phenotype, suggesting tissue-specificity in the secretion process. Using *Sec24d* KO mice and cultured cells, we show that SEC24A and SEC24B also contribute to ER export of procollagen. In contrast, fibronectin 1 requires either SEC24C or SEC24D for ER export. On the basis of our results, we propose that procollagen interacts with multiple SEC24 paralogs for efficient export from the ER, and that this is the basis for tissue-specific phenotypes resulting from SEC24 paralog deficiency.

## INTRODUCTION

Collagens comprise about a third of the total protein in humans and are the major constituent of the extracellular matrix (ECM)(1). Collagens are synthesized as pre-procollagen in the endoplasmic reticulum (ER) and exit the ER as procollagen, which is rapidly processed to a mature form in the Golgi and in the extracellular space of animal tissues (1, 2). The exit of procollagen from the ER depends on the coat protein complex II (COPII) (3, 4). SEC24 proteins are COPII components that are mainly responsible for sorting cargo molecules at ER exit sites (5). There are four SEC24 paralogs (SEC24A-D) in vertebrates and each SEC24 paralog has multiple cargo binding pockets, partly accounting for the ability of COPII to recognize diverse cargo molecules (6-9). The SEC24 paralogs are classified into two subgroups based on their primary amino acid sequences and structures: one includes SEC24A and SEC24B, the other includes SEC24C and SEC24D. These two subgroups each display shared and distinctive cargo selectivity, mediated by selective binding of various transport signals on cargos and SNARE proteins (7, 10, 11).

*SEC24* mutations tend to cause tissue-specific defects, depending on the nature of the mutation and on the organism. *Sec24a* knockout (KO) mice develop and live normally except that mutant animals exhibit markedly reduced plasma cholesterol levels due to defective ER export of PCSK9, a negative regulator of the low-density lipoprotein receptor (12). *Sec24b* deletion causes a completely open neural tube in mice, which occurs because VANGL2 fails to be sorted into COPII vesicles (13). *Sec24c* disruption results in embryonic lethality (14). Conditional knockout of *Sec24c* in neural progenitors during embryogenesis causes apoptotic cell death of post-mitotic neurons in the cerebral cortex (15). The neuronal cell death caused by *Sec24c* ablation was rescued in knock-in mice expressing *Sec24d* in place of *Sec24c*, which indicates a functional equivalency of SEC24C and SEC24D for this aspect (15). Compound heterozygous *SEC24D* mutations cause osteogenesis imperfecta with craniofacial dysplasia in humans (16). In mice, homozygous *Sec24d* deletion (*Sec24d*^*gt/gt*^, a gene-trap mutation) cause embryonic lethality near the 8-cell stage and hypomorphic *Sec24d* KO mutants (*Sec24d*^*gt2/gt2*^) survive to mid-embryogenesis (embryonic day, E13.5) (17). Because of the early embryonic lethality, the role of SEC24D in skeletal formation remains unknown in mice. *sec24d* mutant fish display severe craniofacial and skeletal dysplasia (18, 19). The cartilage cells of the *sec24d* mutant fish accumulate procollagen in the ER, which is consistent with the observed skeletal defects. *sec24c* morphants do not show craniofacial cartilage dysmorphology (18). However, craniofacial development in *sec24c;sec24d* double morphants arrested earlier than in *sec24d* single mutants, indicating that Sec24C has a partial role in chondrogenesis. Thus, among SEC24 paralogs, SEC24D is the most critical for skeletal formation both in humans and in zebrafish. These *in vivo* observations suggest that SEC24D and, to a lesser degree, SEC24C are involved in ER export of procollagens.

The severe bone fragility in patients carrying *SEC24D* mutations and the strong accumulation of procollagen in the ER in *sec24d* mutant fish predict that collagen secretion should be impaired in human SEC24D patients’ cells. Skin fibroblasts derived from a SEC24D-deficient patient, however, displayed only a mild accumulation of collagen in the ER (16). Thus, there seems to be other cellular factors that modulate the collagen secretion phenotype in SEC24D-compromised cells and tissues. In this study, we present evidence that other SEC24 paralogs are the cellular factors compensating for the SEC24D deficiency.

## RESULTS

### The processing of procollagen depends on SEC24D and other factor(s)

To examine the involvement of SEC24D in collagen secretion, we generated *Sec24d*^*gt2/gt2*^ mice as previously described (17). As the *Sec24d* KO mice are embryonically lethal near E13.5 (prior to skeletal development) we decided to monitor collagen processing in embryonic tissues. Procollagen is proteolytically processed to collagen in post ER compartments as procollagen peptidases are present both in the Golgi and in the extracellular space (Fig. 1A) (2). ER export inhibition imposed by *Sec23a* deletion results in pronounced ER accumulation of procollagen and an almost complete inhibition of the proteolytic processing of procollagen *in vivo* (20). Thus, we anticipated consistent, similarly severe collagen phenotypes in the *Sec24d*^*gt2/gt2*^ embryos. Surprisingly, however, the mutant embryos showed a variable and relatively modest processing defect of alpha-1 type I procollagen, pro-α1(I) (Fig. 1B, compare 36d KO with 36e KO). Interestingly, we observed a prominent inhibition of procollagen processing to the mature form (Fig. 1C, see 92a and 92d) and occasionally aberrant processing (Fig. 1C, see 92a, open arrowhead) in the yolk sacs of *Sec24d* KO mice. These results suggest that SEC24D plays an important role in the ER export of procollagen, but that a tissue-dependent factor(s) influences the necessity for SEC24D.

**Figure 1.**
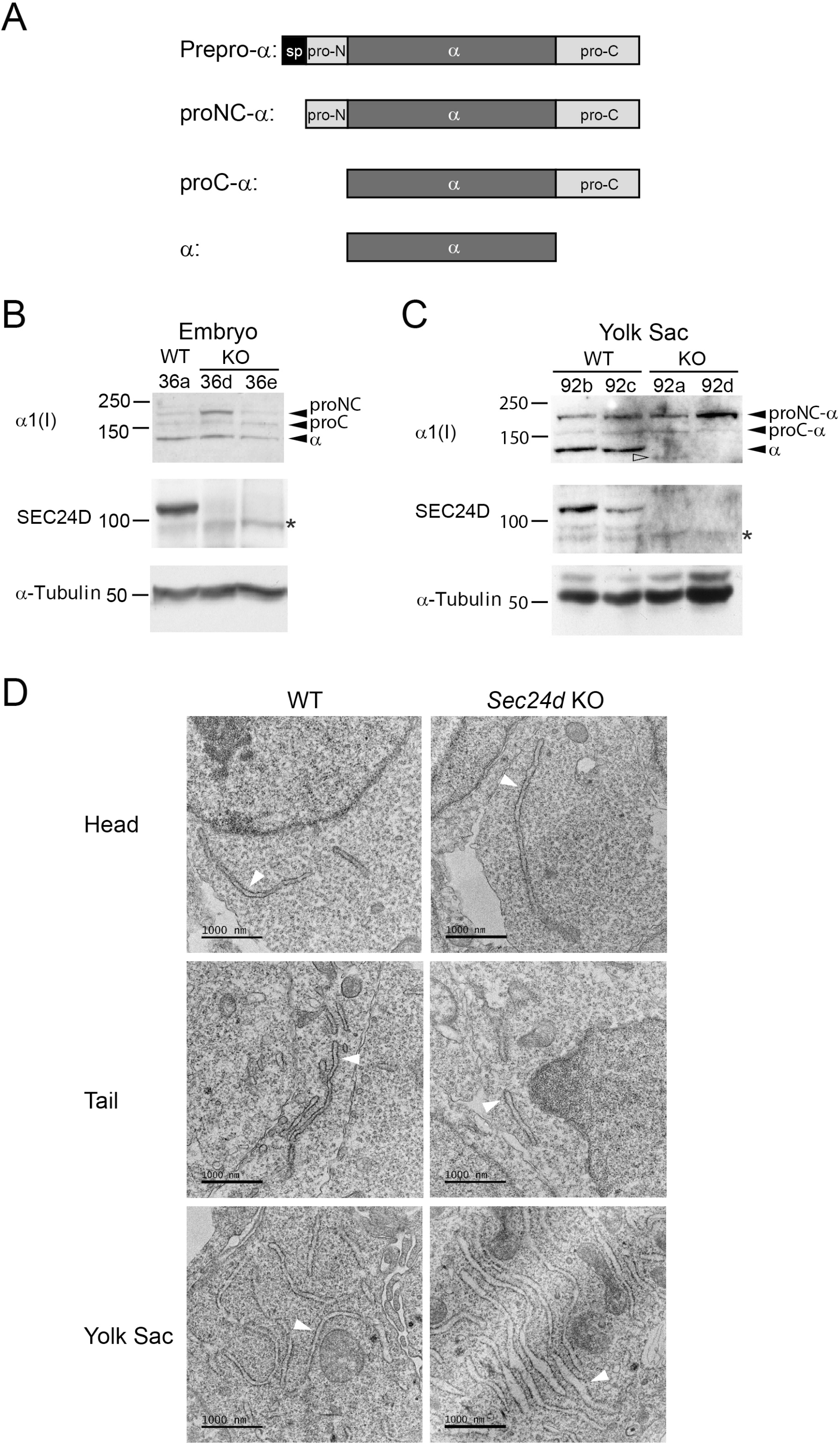
Tissue-dependent phenotypes in *Sec24d* KO tissues. (A) A schematic illustration of collagen processing. Extracts of embryos (B) and yolk sacs (C) were resolved and analyzed with immunoblotting. Asterisks, non-specific bands. Closed arrowheads, normal collagen species. Open arrowhead, abnormal collagen species. α-tubulin was used as a loading control. (D) Embryos and yolk sacs were processed for TEM. A part of the head, the tail and the yolk sac of wild-type and *Sec24d* KO mice are shown. ER cisternae were indicated by white arrowheads.

### Collagen accumulates in the ER of the yolk sacs of Sec24d KO mice

We performed immunohistochemistry on embryonic tissues to visualize collagen distribution patterns and potential disruptions. Given the observed mild accumulation of collagen in the ER of skin fibroblasts from a SEC24D patient (16), we examined collagen signals in the embryonic head skin. Wild-type and *Sec24d* KO embryos had similar distribution of α1(I) within head skin, with most fibroblasts displaying consistent punctate signal throughout the tissue and some deposition distal to superficial cells (Fig. 2A-D). Sporadic cells with more intense α1(I) signal appeared in both genotypes.

**Figure 2.**
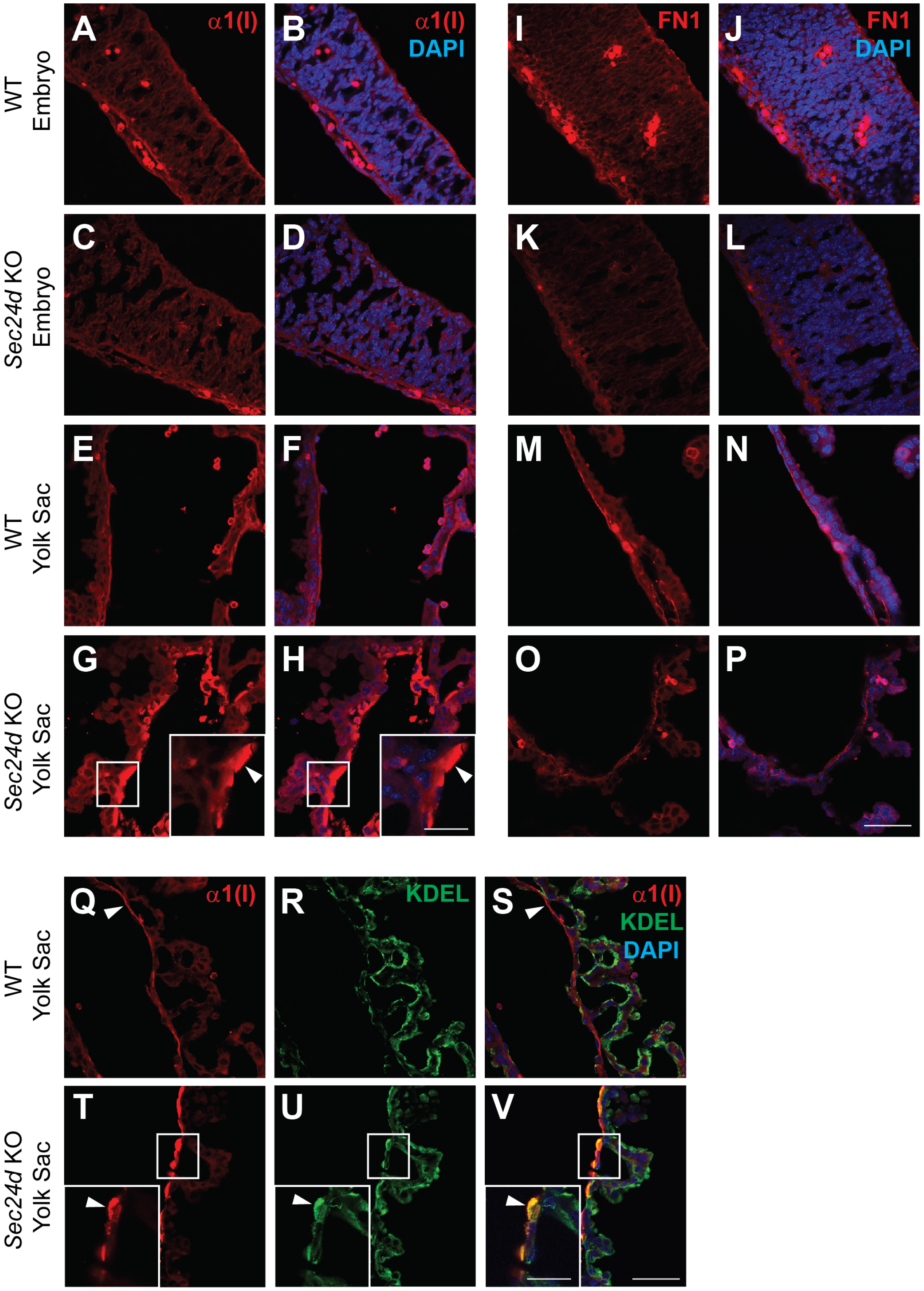
Collagen accumulates in the ER of yolk sacs of *Sec24d* KO mutants. (A-H) Anti-α1(I) immunoreactivity in WT (A-B, E-F) and *Sec24d* KO mutant (C-D, G-H) head skin (A-D) and yolk sac (E-H). Embryo skin displayed similar immunoreactivity patterns in both genotypes. In the yolk sac, *Sec24d* KO mutants had an apparent massive accumulation of pro-α1(I) in the cells lining the lumen (white arrowheads). FN1 expression patterns were similar between genotypes in both the embryo head skin (I-L) and in the yolk sac (M-P). Insets in the bottom right of G and H represent higher magnification images of the boxed regions. Scale bar in H inset is 20 µm for both insets. Scale bar in P is 50 µm for all other images. (Q-V) Anti-α1(I) immunoreactivity in WT (Q, S) and *Sec24d* KO mutant (T, V) yolk sac. Anti-KDEL immunoreactivity in WT (R, S) and *Sec24d* KO mutant (U, V) yolk sac.

When comparing patterns within the yolk sacs, however, striking differences were apparent. The inner mesoderm layer of the wild-type yolk sac has a high collagen content in the ECM (21). We also probed for distribution of type IV collagen. However, type IV collagens were found inside the cell in both genotypes (Fig. S1), preventing us from evaluating its secretion defect. Wild-type tissues displayed diffuse α1(I) signals throughout the yolk sac cells, with distinct, thin ECM deposits on the luminal side of the inner mesoderm cells (Fig. 2E, F, Q, and S). *Sec24d* KO yolk sacs lacked these deposits, instead displaying extensive ER accumulation (Fig. 2G, H, T, and V). The luminal mesoderm cells were distended and malformed, with α1(I) deposits occupying a large portion of the cell body. We also examined the localization of another secretory protein of the ECM, fibronectin 1 (FN1), in both tissues but did not observe any differences between the genotypes (Fig. 2I-P). Together, these results support a cargo-specific, tissue-specific necessity for SEC24D in ER export.

### The factor(s) affecting the tissue dependency is likely a component involved in ER export

An ER export block can enlarge ER cisternae. In particular, the ER accumulation of procollagen can contribute to ER enlargement as collagens are the most abundant secretory proteins (1). Thus, we expected to observe a significant ER distension in the tissues of *Sec24d* KO mice. Unexpectedly, however, almost no ER distension was observed in the cells of the head or the tail in the mutants (Fig. 1D). A significant ER distension was observed in the yolk sac in the mutants. These results are in line with our biochemical and immunohistochemical results showing prominent collagen processing defects in the mutant yolk sacs. Importantly, this ER distention is not fully explained by a simple lack of SEC24D, as this phenotype is completely absent in skin fibroblasts. One possible explanation is that some tissues express another component of ER export machinery that is able to compensate for loss of SEC24D. Thus, we hypothesized that the severe phenotypes we observed in the yolk sac are caused by combined deficiencies in SEC24D and at least one other ER export protein that has low level or negligible expression in this tissue.

### SEC24B is deficient in the yolk sac

We suspected that the unknown tissue-specific compensatory factor(s) is a SEC24 paralog. For example, SEC24C is close to SEC24D in its amino acid sequence, 3D structure, and cargo preference (7, 10, 11), but its expression levels may differ in the embryo and the yolk sac. To test this possibility, we probed for the expression of *Sec24* paralogs in the extracts derived from embryos and yolk sacs (Fig. 3). We measured *Sec24a* transcript levels using a real-time quantitative PCR (RT-qPCR), because we were not able to obtain a reliable antibody specific for mouse SEC24A. *Sec24a* transcript levels were significantly higher in yolk sacs than in embryos (Fig. 3A, WT), while levels of SEC24D were comparable between the two (Fig. 3B, see WT). Interestingly, the levels of SEC24B and SEC24C were very low in wild-type yolk sacs (Fig. 3B, see WT), but collagen processing proceeded normally in this tissue (Fig. 3C, WT yolk sacs). As we examined more WT yolk sacs, we noticed significant individual variations in SEC24C levels (Fig. S2). However, the levels of SEC24B were low consistently. Importantly, the processing of pro-α1(I) occurred normally in wild-type yolk sacs (Fig. 3C and Fig. S2). This result suggests that both SEC24B and SEC24C can be dispensable and that SEC24A and SEC24D can be sufficient to export procollagen efficiently from the ER at least in yolk sacs.

**Figure 3.**
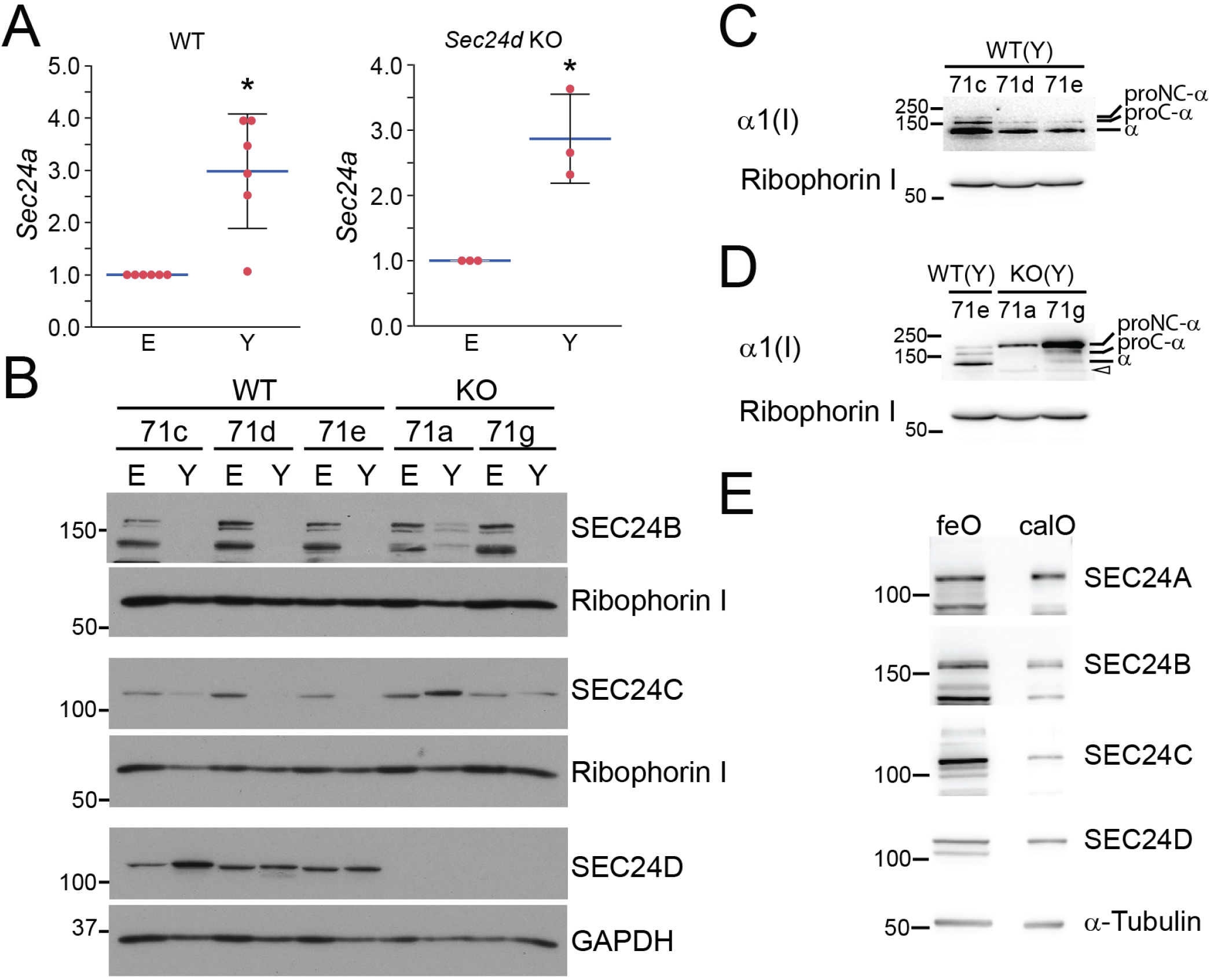
SEC24B is deficient in the yolk sacs of *Sec24d* KO mutants. (A) Levels of *Sec24a* mRNAs were measured with RT-qPCR and were normalized to those of *Gapdh* mRNAs. E, embryo; Y, yolk sac. (A) Student’s t-test: WT, P*<0.005, n=6; *Sec24d* KO, P*<0.01, n=3. Error bars represent standard deviation. (B) SEC24B, SEC24C, and SEC24D were probed with immunoblotting using the extracts of wild-type (WT) and *Sec24d* KO (KO) mutant animals. E, embryo; Y, yolk sac. Ribophorin I or GAPDH was used as a loading control. (C and D) α1(I) was probed in the yolk sac extracts of WT (71c, 71d, and 71e) and *Sec24d* KO mutants (71a and 71g). Open arrowhead, abnormally processed collagen (see Fig 1C 92a animal). Ribophorin I was used as a loading control. (E) SEC24s were probed in lysates from human femoral osteoblasts (feO) and human calvarial osteoblasts (calO).

We also probed for the expression of *Sec24* paralogs in the mutant yolk sacs. Levels of *Sec24a* transcripts in *Sec24d* KO yolk sacs were comparable to those in wild-type yolk sacs (Fig. 3A). As observed in WT yolk sacs, we observed significant variations in the levels of SEC24C, but not in those of SEC24B in mutant yolk sacs (Fig. 3B and Fig. S3). However, higher levels of SEC24C did not affect collagen processing in the mutants (see 71a and 71g in Fig. 3B and D). It is not clear why there were far lower levels of procollagen in the 71a animal than in the 71g animal (Fig. 3D).

Interestingly, we occasionally observed a cellular accumulation of FN1 in *Sec24d* KO yolk sacs (Fig. S3C and D). This could be due to an occasional low expression of *Sec24c* (Fig. S3A) because both SEC24C and SEC24D are lacking under this condition in *Sec24d* KO yolk sac.

We then evaluated the relative levels of SEC24 paralogs in human bone-related cell lysates (Fig. 3E). Interestingly, calvarial osteoblasts showed reduced levels of SEC24B and SEC24C, compared to femoral osteoblasts. It is important to note that the ossification of long bones occurs through chondrocytes and that of neurocranium occurs through calvarial osteoblasts (22). Unfortunately, we were not able to obtain femoral (epiphyseal) chondrocytes. However, the two human bone cell lysates clearly show differential expression of *SEC24* paralogs. It is thus likely the low levels of SEC24B and especially SEC24C in calvarial osteoblasts could account for the severe skull phenotypes documented in SEC24D patients (16). It remains to be seen whether epiphyseal chondrocytes exhibit low expression levels of *SEC24* paralogs, but if this were the case it could potentially account for the severe fragility of the lone bones in SEC24D patients.

### Knock down (KD) of an individual SEC24 paralog has a minimal effect on collagen export

The analyses of the mouse tissues suggest that the absence of SEC24D leads to a mild to modest reduction in ER export of procollagen in embryos. Interestingly, both SEC24B and SEC24C are dispensable for procollagen ER export as observed in wild-type yolk sacs. However, this does not necessarily mean that SEC24B and SEC24C do not contribute to the process. This is partly because the absence of SEC24B appears to contribute to the stronger block of procollagen ER export in mutant yolk sacs. Thus, we decided to systematically test the involvement of SEC24 paralogs in collagen secretion.

To test for the role of SEC24s in collagen secretion, we knocked down the expression of *Sec24* paralogs using RNA interference (RNAi) in wild-type mouse embryonic fibroblasts (MEFs). Our RNAi approach yielded about 70% depletion of SEC24s (Fig. S4). It should be noted that the proteolytic processing of the precursor forms of collagen is inefficient in cultured cells (2) as the collagen processing proteases are diluted in culture medium, resulting in the presence of procollagen in the culture medium (Fig. S5A). In addition, the culture medium used in our experiments did not contain any detectable α1(I) (Fig. S5B). As a control of a secretory cargo, we also monitored FN1 (Fig. S5C). We reasoned that knockdown (KD) of specific SEC24 paralogs would result in an accumulation of secretory proteins in the ER and a reduction of the secretory proteins in the culture medium. We did not observe such effects with the KD of individual SEC24s (Fig. 4A and B). Consistent with this result, collagen did not accumulate in *Sec24d* KO MEFs (Fig. S6), suggesting that SEC24 paralogs have a redundant function for ER export of procollagen and FN1.

**Figure 4.**
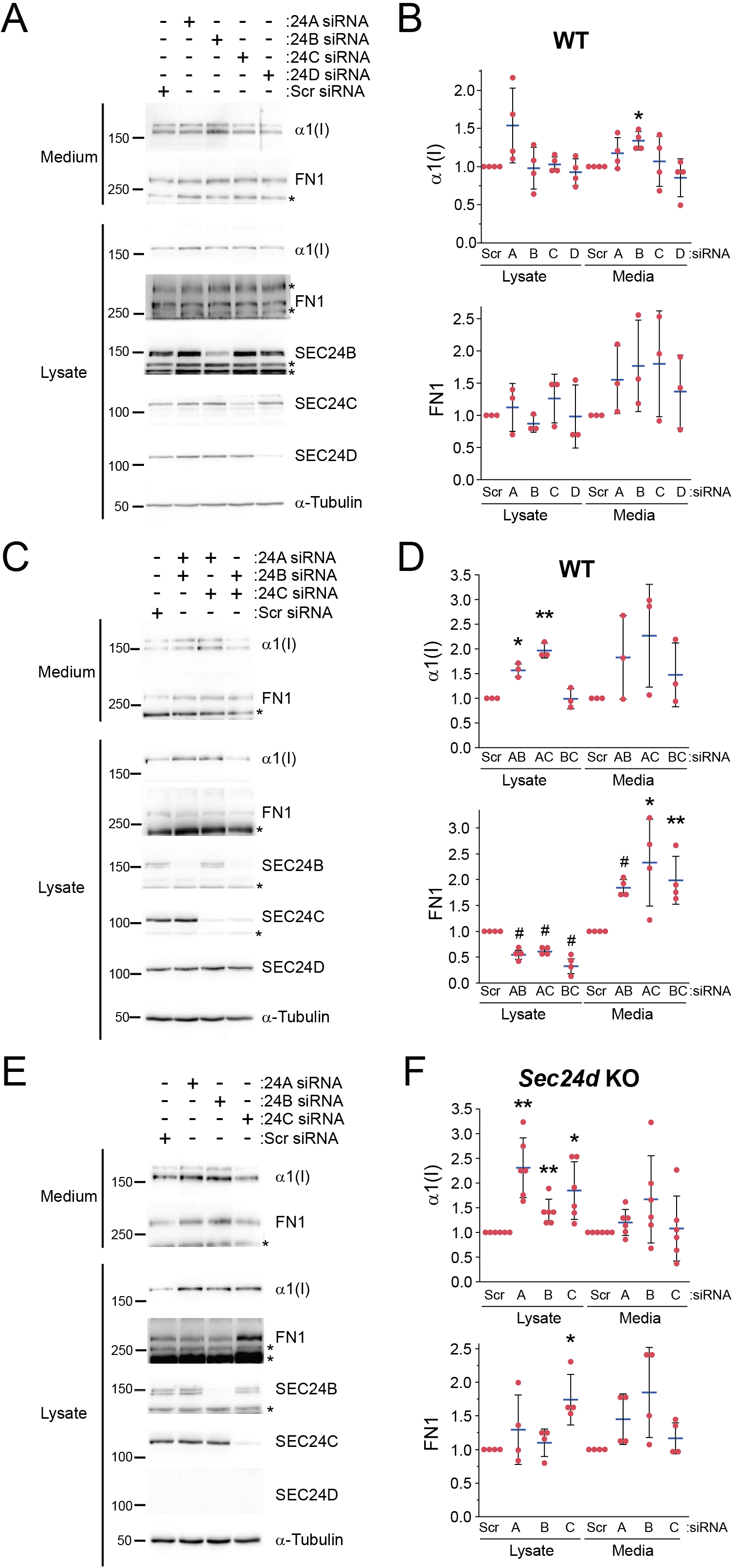
SEC24B and SEC24C have the least effect on collagen secretion. Wild-type (A-D) or *Sec24d* KO (E and F) MEFs were transfected with indicated siRNAs (50 nM) at day 0. The culture medium was replaced with fresh medium at day 1. Cells and a conditioned medium were harvested at day 2. Cell lysates and media were analyzed by immunoblotting (A, C and E). The levels of proteins in the lysates were normalized to those of α-tubulin and the levels of α1(I) and FN1 in the medium were normalized to total protein amounts. In each experimental set, the levels of α1(I) and FN1 were normalized to those in control (scrambled siRNA). Asterisks, nonspecific proteins. Scr, scrambled. Statistical significance was determined with Student’s t-test. Error bars represent standard deviation and middle lines represent average. (B) A KD of *Sec24b* led to an increase of secreted collagen, which happened to coincide with the increased expression of *Sec24a* (Fig. S3). P*<0.05, n=3-4. (D) α1(I): P*<0.05, P**<0.0005, n=3; FN1: P*<0.05, P**<0.01, P^#^<0.0001, n=4. (F) α1(I): P*<0.01, P**<0.005, n=6; FN1: P*<0.01, n=4.

### SEC24A and SEC24D play a major role in procollagen export in MEFs

We then knocked down the expression of two *Sec24* paralogs simultaneously (Fig. 4C-F and Fig. S7A and B). When we attempted to deplete two SEC24 paralogs we noticed that SEC24D KD became ineffective for unknown reasons. Perhaps SEC24D siRNAs were outcompeted by other SEC24 siRNAs for the RNA-induced silencing complex (23). To circumvent this problem, we used *Sec24d* KO MEFs when SEC24D depletion is needed. Knocking down SEC24B/C did not affect collagen levels (Fig. 4C and D), which was consistent with the mouse tissue results (Fig. 3B and C, WT). Interestingly, a simultaneous KD of SEC24A with any SEC24 paralog resulted in cellular accumulation of collagen (Fig. 4C-F). Similar results were obtained with a KD of SEC24D with any SEC24 paralog (Fig. 4C-F). These results underscore the importance of SEC24A and SEC24D as compared to SEC24B and SEC24C. In contrast, FN1 accumulated in the cell only when SEC24C/D were knocked down (Fig. 4C-F), suggesting that either SEC24C or SEC24D can export FN1 efficiently from the ER. It is important to note that collagen is mostly found in the ER even in the control cells (Fig. 6A, Scr), but when the ER export is blocked, the ER collagen signals increase (Fig. 6A, ABC). The levels of secreted collagen and FN1 were not reduced (Fig. 4C-F). This modest collagen phenotype likely occurred because we could not inhibit a target gene expression completely with RNAi.

To test if the increased cellular collagen levels result from collagen expression, we probed *Col1a1* mRNA levels with RT-qPCR (Fig. S8). Although there were instances where *Col1a1* mRNA levels increased, it was increased slightly or not observed consistently. To exclude any contribution from *Col1a1* gene expression, we performed cycloheximide chase experiments in wild-type MEFs. Cellular collagen was cleared slower under SEC24A/B KD or SEC24A/C KD condition than control (Scr) or SEC24B/C KD condition (Fig. S9A and B). To inhibit lysosomal degradation of collagen, we also performed cycloheximide chase experiments in the presence of bafilomycin A1. We observed slower clearance of collagen under SEC24A/B KD or SEC24A/C KD condition than control (Scr) or SEC24B/C KD condition (Fig. S9C and D). These results are consistent with delayed ER export of procollagen.

We also measured the expression levels of genes relevant to procollagen ER export (Fig. S8). Interestingly, *Sedlin* expression was upregulated in KO MEFs when additional SEC24 was knocked down (Fig. S8, KO). In addition, *Sec23b, Sedlin*, and *klhl12* were upregulated under SEC24B/C KD condition (Fig. S8, WT). Perhaps these upregulations may partly account for the absence of collagen phenotype in SEC24B/C KD condition. However, in spite of similar upregulation (Fig. S8, KO), we observed a significant cellular collagen accumulation in SEC24C/D KD condition (Fig. 4E and F). Such upregulation was not observed when all SEC24s were knocked down (Fig. S10). Thus, the contribution of the upregulated genes to procollagen ER export is unclear in our experimental conditions. We also tested if dysregulation in the unfolded protein response (UPR) of the ER imposed by SEC24 KD contributed to the increased collagen levels especially in double KD conditions. UPR markers tested were largely unaffected (Fig. S11).

The observation that A KD of two SEC24 paralogs did not reduce the levels of secreted protein likely reflects a modest ER export defect. As a control, we tested whether our RNAi approach could allow us to block collagen secretion. A KD of three SEC24 paralogs also resulted in relatively modest collagen phenotypes such as an increase in cellular collagen levels or a modest decrease in secreted collagen levels (Fig. 5A-D and Fig. S7C and D). Interestingly, in the case of FN1, a KD of SEC24A/B/C or SEC24A/B/D did not lead to an increase in cellular FN1 levels. This result is consistent with the idea that SEC24C or SEC24D alone can support an efficient ER export of FN1.

**Figure 5.**
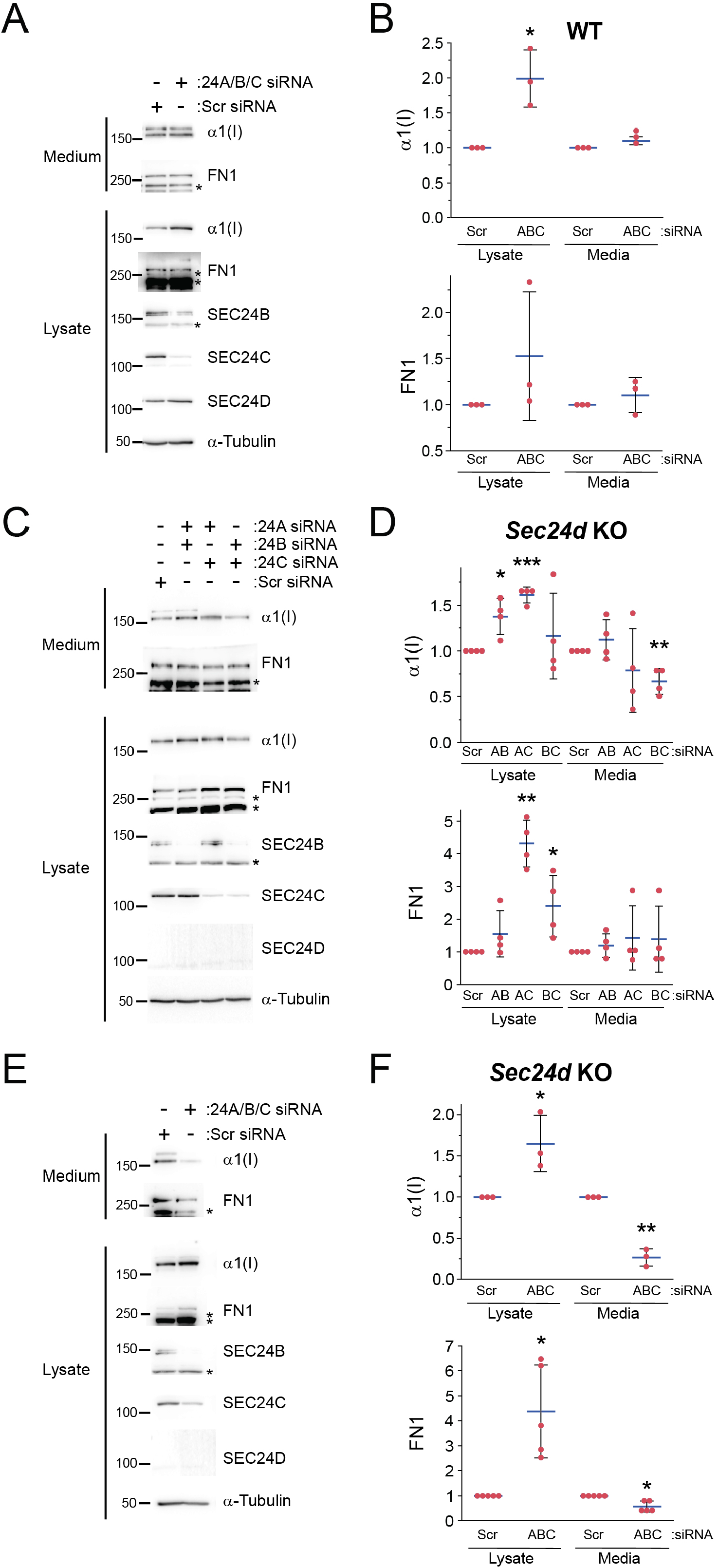
All SEC24s contribute to collagen secretion. Wild-type (A and B) or *Sec24d* KO (C-F) MEFs were treated and analyzed as described in the legend of Fig. 4. Asterisks, nonspecific proteins. Scr, scrambled. Statistical significance was determined with Student’s t-test. Error bars represent standard deviation and middle lines represent average. (B) P*<0.05, n=3. (D) α1(I): P*<0.01, P**<0.005, P*** <0.0001, n=4. FN1: P*<0.05, P**<0.0001, n=4. (F) α1(I): P*<0.05, P**<0.0005, n=3. FN1: P*<0.05, n=5.

Then, we knocked down the expression of all SEC24 paralogs (Fig. S7E). When the expression of all SEC24 paralogs was reduced, a strong secretion block was achieved (Fig. 5E and F). The levels of *Col1a1* mRNAs were not increased under the condition (Fig. S10), indicating that the cellular collagen accumulation is not the consequence of an increased expression of *Col1a1* mRNAs. Consistent with this collagen secretion phenotype, intracellular collagen was found in the ER (Fig. 6A and B) and the ER cisternae were enlarged drastically under this condition (Fig. 6C). Interestingly, SEC16A signals that represent ER exit sites were enhanced in regions where PDI signals were strong (Fig. S12). These ER regions appeared to be dilated. Morphologies of ERGICs (ERGIC53) and Golgi apparatus (GM130) were unaffected (Fig. S13). The distribution of SEC22B seemed to be shifted from ERGICs to the ER (Fig. S14). It is noteworthy that SEC22B has a SEC24A/B-specific ER export signal whereas ERGIC53 has a C-terminal dihydrophobic ER export signal that can be recognized by all SEC24s (11, 24). When all SEC24s were knocked down, ERGIC53 can use any residual SEC24s for ER export whereas SEC22B can use residual SEC24A and SEC24B only. This likely accounts for the shift of SEC22B to the ER.

**Figure 6.**
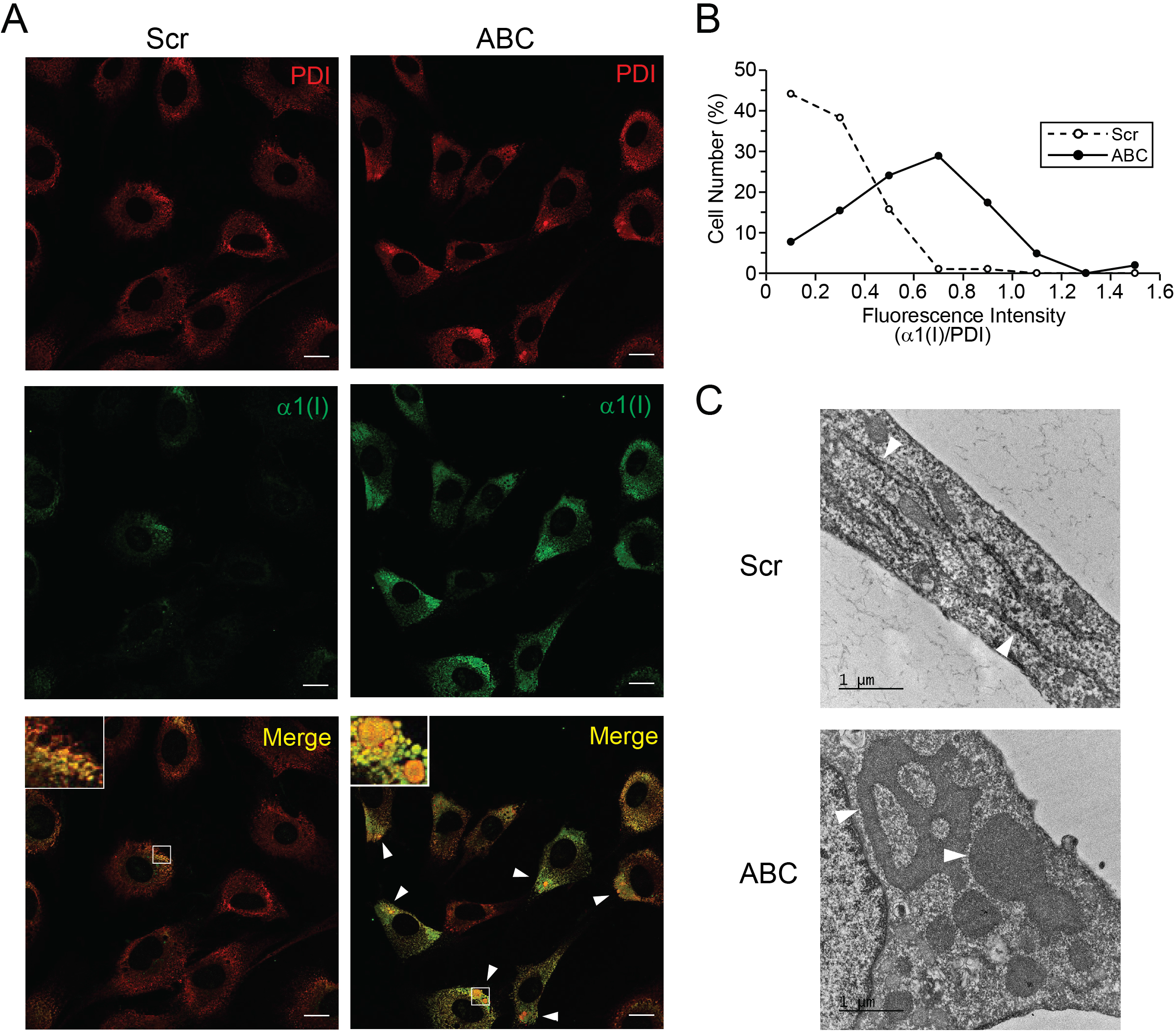
Collagen accumulates in the ER in SEC24-depleted cells. *Sec24d* KO MEFs were transfected with indicated siRNAs (50 nM) at day 0. The culture medium was replaced with fresh medium at day 1. Cells were fixed at day 2 and processed for immunfluorescent labeling (A) or for electron microscopy (C). (A) Protein disulfide isomerase (PDI) was used as an ER marker. Scale bar, 20 μm. Inset, an enlarged image of the indicated region with a box. Arrowheads, a region of the ER that shows strong collagen signals. PDI signals were also stronger in the Sec24-depleted cells, reflecting an increase of the unfolded protein response of the ER. (B) Fluorescent signals (corrected total cell fluorescence) of collagen and PDI were quantified with ImageJ and collagen signals were normalized to PDI signals. Statistical significance was determined with Student’s t-test. P*<0.0001, n=102 (Scr, Scrambled siRNA), n=104 (ABC, siRNAs for *Sec24a, Sec24b* and *Sec24c*). (C) Electron micrographs of *Sec24d* KO MEFs transfected with scrambled siRNA (Scr) or siRNAs for *Sec24a, Sec24b* and *Sec24c* (ABC). ER cisternae were indicated with arrowheads. Scare bar, 1 μm.

We generated a recombinant human *GFP*-*SEC24D* construct under a Tet-on promoter and introduced it into *Sec24d* KO MEFs to rescue the collagen secretion defects. Unfortunately, the expression of the recombinant *SEC24D* was much lower than that of endogenous *Sec24d*, preventing us from properly evaluating the impact of the recombinant gene on collagen in MEFs (Fig. S15A). We then stably expressed the recombinant gene in human U-2 osteosarcoma (U2OS) cells and knocked down all SEC24 expression with siRNAs. KD of SEC24s results a mild collagen phenotype (increase of cellular collagen without corresponding decrease in secreted collagen) (Fig. S15B and C), probably reflecting a mild ER export block. However, when the recombinant *SEC24D* was overexpressed, the cellular collagen levels reverted to normal levels, indicating that such collagen phenotype was caused by SEC24 deficiency, rather than by an off-target effect.

Taken together, our data show that while SEC24A and SEC24D play a major role in ER export of procollagen, SEC24B and SEC24C also contribute to the process, particularly when SEC24A and/or SEC24D are lacking. In the case of FN1, SEC24C and SEC24D act on FN1 redundantly.

## DISCUSSION

Considering the similarities and differences in primary amino acid sequences and cargo binding sites of SEC24s, it is evident how SEC24 paralogs can have both distinct and overlapping cargo preferences. For example, VANGL2 is recognized preferably by SEC24B (13), SEC22 SNARE by SEC24A or SEC24B (Mancias and Goldberg, 2007), STX5 by SEC24C or SEC24D (Mancias and Goldberg, 2008), and p24 and ERGIC53 by any of SEC24s (11, 25). Here we showed that FN1 is favored by either SEC24C or SEC24D (Fig. 5A-D). However, procollagen is unique in that it can utilize SEC24A/D (including SEC24A/C/D), SEC24A/B/C, or SEC24B/C/D for ER export, suggesting that procollagen needs multiple SEC24 paralogs from the two subgroups for efficient ER exit. It is possible that procollagen needs two different cargo adaptors each of which is recognized by a different SEC24 paralog from the two subgroups. Alternatively, procollagen could utilize one cargo adaptor that has two export signals, which are recognized by two SEC24 paralogs. It is also possible that SEC24C/D recognizes a cargo adaptor and SEC24A/B recognizes a molecule that should be co-packaged with procollagen. The exact mechanism remains to be determined.

TANGO1/MIA3 and cTAGE5 (a splice variant of MIA2) bind each other and are believed to function as a cargo adaptor for procollagen (26, 27). The luminal the Src homology 3 (SH3) domain of TANGO1/MIA3 binds type VII procollagen and the cytosolic proline-rich domain binds SEC23 (26-28). TANGO1 is necessary for the secretion of a type IV collagen, Viking, in *Droshophila* (29) and numerous collagens, including type I, II, III, IV, VII and IX collagens in mice and humans (30, 31). Despite the compelling *in vivo* evidence of TANGO1 for the secretion of various collagens, TANGO1 does not appear to bind type I procollagen *in vitro* (26, 27). It has been suggested that HSP47 binds to various types of procollagens and directs them to the SH3 domain of TANGO1 (32). Alternatively, the collagen I secretion defect in TANGO1 KO MEFs is a result of indirect effects (33). Consistent with this possibility, a general role for TANGO1 has recently been shown in maintaining secretory pathway structure and function (34). Thus, it is possible that additional cargo adaptors which are recognized by SEC24A/B contribute to collagen recognition. Consistent with this idea, there have been found candidates of collagen adaptors (35, 36).

As ER-to-Golgi SNAREs are differentially packaged by SEC24A/B and SEC24C/D into COPII vesicles (7, 37), a simultaneous depletion of two SEC24 paralogs within the same subgroup can block the packaging of the SNAREs, which can inhibit procollagen export indirectly. For example, SEC24A/B packages the R-SNARE SEC22B and SEC24C/D packages the Q-SNARE complex (STX5/Sed5p, membrin/Bos1p/GS27 and BET1) into COPII vesicles (7, 10, 25). Thus, a depletion of SEC24C/D or a depletion of SEC24A/B could block procollagen export indirectly because such depletion would block the packaging of the SNAREs into COPII vesicles. However, other combinations (e.g., a depletion of SEC24A/D or that of SEC24B/D) would not interfere with the packaging of the SNAREs into COPII vesicles. In addition, under our *in vitro* conditions, the depletion is not complete. Thus, such indirect effects are unlikely. In fact, a KD of SEC24A/B did not lead to a cellular accumulation of FN1 (Fig. 4C and D).

We observed a substantial secretion block for both collagen and FN1 only when three SEC24 paralogs were knocked down in *Sec24d* KO MEFs. We propose a multi-SEC24 model for efficient procollagen ER export. This model implies that two distinct ER export signals are embedded in two different adaptors and each of the ER export signals is recognized by SEC24A/B and SEC24C/D, respectively. However, our *in vitro* observations, in particular, can be explained by a threshold effect model, namely that a threshold level of SEC24s, regardless of paralogs, is required for efficient ER export. This model implies that collagen preference by each SEC24 subgroup is similar so that as long as a threshold level of SEC24s is present procollagen can exit the ER efficiently. We observed that SEC24 requirements for procollagen and FN1 are different. Perhaps, the threshold level of SEC24s that is required for procollagen ER export is different from that for FN1. Although two SEC24 subgroups show distinct cargo preference, they do have comparable cargo preference for the C-terminal dihydrophobic ER export signal (11). The threshold model likely involves such a shared export signal. Interestingly, a mutation (S1015F) in a SEC24C/D-specific cargo binding site was identified from a human SEC24D patient (16). Although this site recognizes the IxM motif of syntaxin 5 (7, 25), it is not clear if this site also recognizes a procollagen adaptor. Thus, questions remains about cargo-binding sites of SEC24 and cargo adaptors necessary procollagen ER export.

The vast majority of yolk sacs of *Sec23a* KO mutants are broken probably due to a severe collagen secretion block in the mutant yolk sacs (20). We have not observed any broken yolk sacs in our mutants. Perhaps the secretion block in our mutant yolks sacs is not as severe as that in the *Sec23a* mutant because SEC24A and SEC24C can contribute to collagen secretion to a certain degree. In fact, FN1 was secreted normally (Fig. 2). Alternately, the yolk sac breakage may also involve other secretory proteins that are secreted normally in *Sec24d* KO yolk sacs. *Sec23a* KO mutants also present with reopening of the closed neural tube (20). Such phenotype was not observed in our mutants, nor in the previous work describing *Sec24d* KO mutants (17). This reopening was caused by a collagen secretion defect in skin fibroblasts, which likely compromises the tensile strength of the skin tissue. We observed only a modest processing defect of collagen in embryo extracts (Fig. 1B). In addition, the ER of the head cells was not distended in the mutants (Fig. 1D). Thus, we do not expect a reduction in the tensile strength of the skin tissue of our mutants. Together, our work demonstrates that the unique tissue-specific consequences of *Sec24d* KO result from the variable presence of other SEC24 paralogs, and that SEC24 paralogs cooperate in a tissue-specific and cargo-specific manner to facilitate ER export of secretory proteins.

## MATERIALS AND METHODS

### Ethics Statement

All animal care and use complied with the Principles of Laboratory and Animal Care established by the National Society for Medical Research. The Institutional Animal Care and Use Committee (IACUC) of the Iowa State University approved all animal protocols in this study under protocol number 19-061. For the immunohistochemical work, all mice were housed in an AAALAC International-accredited facility under IACUC approval (protocol #156-01-22D, Sanford Research, Sioux Falls, SD, USA).

### Generation of *Sec24d* KO mice

ES cell clone RRR785 was obtained from the International Gene Trap Consortium (IGTC, Bay Genomics, San Francisco, CA, RRID:CVCL_NQ03), and was referred to as *Sec24d*^*gt2*^. ES cell culture and microinjection, and recovering the mutant allele were performed as described previously (17). Mice from *Sec24d*^*gt2*^ (RRID: 5515787) were genotyped using a PCR. A PCR reaction including a common forward primer (In20F1) (Table S1) located upstream of the insertion site in intron 20 and a reverse primer located downstream of the insertion site (In20R1) in intron 20 produces a 715 bp product from the wild-type allele. A PCR reaction including primer In20F1 and a reverse primer V20 located in the gene trap vector produces a 558 bp product from the gene trap allele.

### Timed Mating

Timed matings were performed by intercrossing *Sec24d* heterozygous mice. Copulation plugs were noted next morning and counted as day E0.5 of embryonic development. Embryos were harvested from E8.5–E13.5 for genotyping and E12.5 for the preparation of mouse embryonic fibroblasts, tissue extracts, and histology samples. Genotyping was performed on genomic DNA isolated from yolk sacs or embryos.

### Cell culture and transfection

Wild-type and mutant mouse embryonic fibroblasts (MEFs) were obtained from wild-type and knockout mice, respectively. All cell lines were maintained in Gibco™ high glucose DMEM supplied with 10% FBS (Cat. 11995065 and 16000044, Thermo Fisher Scientific, Waltham, MA, USA). Human osteosarcoma cells, U-2 OS, were purchased from American Type Culture Collection (ATCC, Manassas, VA, USA, RRID: CVCL_0042). U2OS cells were maintained in Gibco™ McCoy’s 5A medium with 10% FBS (Cat. Cat. 11995065 and 16000044, Thermo Fisher Scientific, Waltham, MA, USA). A knockdown of a target gene was achieved with transfecting cells with specific siRNA (Table S2, Horizon Discovery, Lafayette, CO, USA) with Lipofectamine RNAiMAX reagent (Cat. 13778, Thermo fisher Scientific, Waltham, MA, USA) according to the manufacturer’s protocol. MEFs were seeded one day prior to transfection. Fifty nano molar of siRNAs (Table S2) and Lipofectamine RNAiMAX were mixed and added to DMEM in the absence of FBS at day 0. Media were removed and replaced with complete DMEM media in the following morning. Next day, cells and media were collected. U2OS cells were seeded one day prior to transfection. Twenty-five nano molar of siRNAs (Table S2) and Lipofectamine RNAiMAX were mixed and added to McCoy’s 5A in the absence of FBS at day Media were removed and replaced with complete McCoy’s 5A media with or without 1 μg/ml doxycycline in the following day. Next day, cells and media were collected. The lysates of human femoral osteoblasts and human calvarial osteoblasts were purchased from ScienCell Research Laboratories (Cat. 4606 and 4656, Carlsbad, CA, USA).

### Plasmids

Human *SEC24D* was cloned from pJK15 plasmid into pAcGFP1-C1 (Addgene, Watertown, MA, USA) to generate a *GFP-SEC24D* fusion gene. *GFP-SEC24D* sequence was amplified with PCR using primers described in Table S1. These PCR products were used to insert *GFP-SEC24D* into a lentivirus vector (pLVX-TRE3G) through In-Fusion Cloning, following manufacturer’s protocol (Takara Bio, Mountain View, CA, USA). Lentivirus vectors pLVX-Tet3G and pLVX-TRE3G (Cat. 631358 and 631193) were purchased from Takara Bio.

### Generation of Tet-on human GFP-SEC24D in U2OS cells

All Lenti-X products and kits were purchased from Takara Bio USA Inc. (Mountain View, CA, USA). To generate lentiviruses, Lenti-X 293 packaging cells (Cat. 632180) were seeded one day prior to transfection. Seven micrograms of lentivirus vectors (pLVX-Tet3G or pLVX-TER3G/GFP-SEC24D) were diluted with 600 μl water and mixed with Lenti-X Packaging Single Shot (Cat. 631275). The mixture was incubated at RT for 10 min. The entire mixture was added to 8 ml of culture media and incubated with the packaging cells overnight. Then, additional 6 ml of culture media were added to the cells and incubated further (48h) to collect virus-containing media. Lentiviral titer was measured with Lenti-X qRT-PCR Titration Kit. U2OS cells were co-infected with viruses harboring pLVX-Tet3G and pLVX-TRE3G-GFP-SEC24D with 1:1 multiplicity of infection ratio in the presence of 4 μg/ml polybrene. Viruses were removed after 24 h of infection. Cells were selected with antibiotic (600 μg/ml G418 and 1 μg/ml puromycin) (Cat. G5013 and Cat. 540222, MilliporeSigma, Burlington, MA, USA) to generate cells stably expressing *GFP-SEC24D* in the presence of doxycycline (1 μg/ml).

### Cycloheximide chase analysis

Cells were seeded one day prior to transfection. Fifty nano molar of siRNAs (Table S2) and Lipofectamine RNAiMAX were mixed and added to DMEM in the absence of FBS. Media were removed and replaced with complete DMEM media in the following day. Next day, cells were treated with fresh media containing cycloheximide (100 μg/ml) and collected at indicated times.

### Immunoblotting and antibodies

Animal tissues were lysed in Triton X-100 buffer (150 mM NaCl, 50 mM Tris-HCl, 1% TritonTM X-100, 0.05% SDS, 1 mM EDTA). Cultured cells were lysed in a RIPA buffer (Cat. 9806s, Cell Signaling, Denvers, MA, USA). Cellular extracts were resolved in a 9% SDS-polyacrylamide gel and transferred to PVDF membranes (Cat. IPVH00010, MilliporeSigma, Burlington, MA, USA) through a wet transfer. Membranes were blocked in 5% skim milk in Tris-buffered saline with 0.1% Tween 20 (TBST) for 1 hour and then incubated with primary antibodies. The membranes were washed with TBST and then incubated secondary antibodies conjugated with horseradish peroxidase. Proteins were visualized with an ECL western blotting detection kit (GE Healthcare, Chicago, IL, USA) through an Azure c600 gel imaging system and bands were quantified with an Azurespot software (Dublin, CA, USA). Anti-SEC24A antibody (Cat. 9678, RRID:AB_10949103) was purchased from Cell Signaling Technology (Danvers, MA, USA). Anti-fibronectin 1 antibody (15613-1-AP) and anti-ERGIC-53 (13364-1-AP, RRID:AB_2135994) were purchased from Proteintech (Rosemont, IL, USA). Anti-SEC24B antibody (A304-876A, RRID:AB_2621071), anti-SEC24C antibody (A304-759A, RRID:AB_2620954), anti-GAPDH antibody (A300-639A) and anti-SEC16A (A300-648A, RRID:AB_519338) were purchased from Bethyl Laboratories (Montgomery, TX, USA). Anti-type I collagen □1 antibody (LF68, ENH018-FP) was purchased from Kerafast Inc.(Boston, MA, USA). For immunohistochemistry, rabbit polyclonal anti-collagen I (203002, MD Bioproducts, Oakdale, MN, USA). Anti-α-tubulin antibody (sc-8035) was purchased form Santa Cruz Biotechnology (Dallas, TX, USA). Anti-SEC24D antibody (#3151) and anti-ribophorin I antibody are gifts from Randy Schekman at UC Berkeley. Anti-mouse PDI antibody (ADI-SPA-891, RRID:AB_10615355) was purchased from Enzolifesciences (Farmingdale, NY, USA). Anti-Sec22b antibody (Cat. 186003) was purchased from Synaptic Systems (Goettingen, Germany). Anti-GM130 antibody (Cat# ab52649, RRID:AB_880266) was purchased from Abcam (Cambridge, MA, USA). For immunohistochemistry, anti-COLA1 antibody was purchased from MD Bioproducts (Cat. 203003, St. Paul, MN, USA) and anti-COLIV antibody (Cat. NB120-6586) and anti-KDEL antibody (NBP1-97469) were obtained from NOVUS (Centennial, CO, USA). The anti-COLIV antibody was raised against bovine placental type IV collagen.

### Immunofluorescence and confocal imaging

Cells were seeded on a coverslip and fixed in 4% paraformaldehyde then permeabilized by incubation with 0.1% TritonTM X-100 (Fisher Scientific, Waltham, MA, USA). After permeabilization, the seeded coverslip was incubated with primary antibodies for 2 h at RT and followed by 1.5 h of a secondary antibody incubation. Mounting medium Mowiol/DAPI/PPD (MilliporeSigma, Burlington, MA, USA) was applied when the coverslip was centered on slides. Anti-human procollagen type 1 antibody (LF68, ENH018) was purchased from Kerafast Inc. (Boston, MA, USA) and anti-PDI antibody (ADI-SPA-891, RRID:AB_10615355) was purchased from Enzo Life Sciences (Farmingdale, NY, USA). Confocal images were taken with a Leica SP5 X (Exton, PA) confocal microscope at the Roy J. Carver High Resolution Microscopy Facility in the Iowa State University.

Mouse E11.5 embryos and yolk sacs were flash frozen upon collection and sectioned using a cryostat. Tissues were post-fixed, blocked in 15% goat serum in TBS with 0.3% Triton X-100 (TBST) and then incubated with the indicated primary antibodies overnight at 4 oC. The tissues were washed with TBS and then incubated with secondary antibodies conjugated with the appropriate fluorophores for two hours at RT. After mounting a coverslip, tissues were imaged using with a Nikon A1R (Melville, NY, RRID:SCR_020317) confocal microscope at the Sanford Research Histology and Imaging Core Facility.

### Electron Microscopy

MEFs were seeded on a six-well plate and transfected with mouse SEC24s siRNA. Transfected MEFs were fixed in fixative solution (1% paraformaldehyde, 3% glutaraldehyde and 1M pH 7.2 cacodylate buffer) for 24-48 hours. Images were taken with a 200kV JEOL 2100 scanning/transmission electron microscope at the Roy J. Carver high resolution microscopy facility in Iowa State University.

### Quantitative PCR

Cellular RNAs were extracted using a TRIzol™ reagent (Cat. 15596026, Thermo Fisher Scientific, Waltham, MA, USA) according to the manufacturer protocol and complementary DNAs were generated using an AccuScript High Fidelity 1^st^ Strand cDNA Synthesis kit (Agilent Technologies, Santa Clara, VA, USA). A quantitative PCR was performed using a QuantStudio™ 3 Real-Time PCR system (Thermo Fisher Scientific, Waltham, MA, USA, RRID:SCR_020238) with an SYBR green supermix fluorescein (Cat. 1708880, Bio-Rad Laboratories, Hercules, CA, USA) for detection. qPCR primers for mouse collagen (*Col1a1*) (38), *Sec24a, Sec23a* (39), *Sec23b* (39), *Tango1* (31), *Klhl12, Sedlin*, and *Gapdh* were obtained from PrimerBank and their sequence information is shown in Table S1.

### Statistical Analysis

We performed Student’s t-test to assess the significance of differences between two groups of data. We also performed power analysis (α=0.05 and power = 80%) to determine the minimal sample size.

**Table 1.** Summary of phenotypes for SEC24 depletion.

| Tissues | Treatments/Read-out | Observations |
| --- | --- | --- |
| Mouse embryos | pro-α1(I) processing | Mild/moderate processing defects in KO animals. |
| Mouse yolk sacs | pro-α1(I) processing | Severe processing defects in KO animals. |
| Mouse heads & tails | EM | Normal ER morphology in WT and KO animals. |
| Mouse yolk sacs | EM | Distended ER morphology in KO animals. |
| Mouse heads | IHC for α1(I) and FN1 | Normal secretion of α1(I) and FN1 in KO animals. |
| Mouse yolk sacs | IHC for α1(I) and FN1 | ER accumulation of pro-α1(I), but not that of FN1 in KO animals. |
| Mouse embryos | SEC24 levels | Even levels of SEC24 paralogs in WT and KO animals. |
| Mouse yolk sacs | SEC24 levels | Consistently lower levels of SEC24B and occasionally lower levels of SEC24C |
| MEFs | *Sec24d* KO | No change in α1(I) secretion. |
| MEFs | KD of individual SEC24 paralogs | No change in the secretion of α1(I) and FN1. |
| MEFs | KD of two SEC24 paralogs | A cellular accumulation of pro-α1(I) except in SEC24B/C KD. A moderate cellular accumulation of FN1 in SEC24C/D KD. |
| MEFs | KD of three SEC24 paralogs | A cellular accumulation of pro-α1(I). A cellular accumulation of FN1 in SEC24C/D KD. |
| MEFs | KD of four SEC24 paralogs | A secretion block and a cellular accumulation of pro-α1(I) and FN1. A strong ER accumulation of pro-α1(I) and a severe distension of ER cisternae. |

## ACKNOWLEDGEMENTS

The authors acknowledge Dr. Marit Nilsen-Hamilton (Iowa State University) for helping to obtain mouse embryonic fibroblasts. Research reported in this publication was supported by the National Institute of General Medical Sciences of the National Institutes of Health under award number R01GM110373, the Sanford Research Imaging Core within the Sanford Research Center for Pediatric Research (NIH P20GM103620), and the Sanford Research Molecular Pathology Core within the Sanford Research Center for Cancer Biology (NIH P20GM103548).

## AUTHOR CONTRIBUTION

CLL and JK designed the study; SB and JK contributed to the construction of mutant mice, JC and JW prepared for mouse tissues, JC, JB, SO, JW collected IF data in mouse tissues; TM collected the data for CHX experiments. CLL collected the rest of the data; CLL and JK wrote the manuscript. All contributed to editing and reviewing the manuscript.

