## Supplemental Information for "Collagen has a unique SEC24 preference for efficient export from the endoplasmic reticulum"

Table S1. Primers used in this study

| Primer | Sequence (5’-3’) |
| --- | --- |
| In20F1 | GGCAGTGGAAGGTGTAAGGA |
| In20R1 | CTGGCCCTGAATTTATTGTGTG |
| V20 | GACCTGGCTCCTATGGGATA |
| *Col1a1* forward | GAGCGGAGAGTACTGGATCG |
| *Col1a1* reverse | GTTCGGGCTGATGTACCAGT |
| *Gapdh* forward | AATGGATTTGGACGCATTGGT |
| *Gapdh* reverse | TTTGCACTGGTACGTGTTGAT |
| Mouse *Sec24a* forward | CCACAAGTGTCATCGAGTCAA |
| Mouse *Sec24a* reverse | AGAACCACCGTAGTTCGACTG |
| Mouse *Sec23a*  forward  Mouse *Sec23a*  reverse  Mouse *Sec23b*  forward  Mouse *Sec23b*  reverse  Mouse *Tango1*  forward  Mouse *Tango1*  reverse  Mouse *Klhl12*  forward  Mouse *Klhl12*  reverse  Mouse *Sedlin*  forward  Mouse *Sedlin*  reverse | TCAAGGGAAATAAAGATTTCAGGA  AGTTTGAGCATCTGCCCAGT  CTTGTGCCCTTGACCAAACT  CTTCAGTTCCCGAGAGGTCTT  AGCCACGGACGGCGTTTCTC  ATCCCCCTGCCAGTTTGTAGTA  TTCGGAAGCCAGCAGTCTCCTA  CCACATACCGTCTCTTGCGAGT  CTGCTCTTGACCTTGTGGACGA  CCATCTTCTTGCCTCACATCATG |

Table S2. siRNAs used in this study

| siRNA | Sequence (5’-3’) |
| --- | --- |
| Mouse *Sec24a* siRNA (Cat #: M-056263-01-0005) | GCUCAUUACUGAUGCUUG |
|  | GAUGCACGCUGACGAGUGU |
|  | GAAUUCAGUUUGCCAGAGU |
|  | AGGAAGGUAUUACGUCAAA |
| Mouse *Sec24b* siRNA (Cat #: M-048898-01-0005) | CUAAUGUGGUGUAUCCUAA |
|  | CCAUAUGGGCAGAUGUUUA |
|  | AAUCAGAGGUCGAGUACCA |
|  | GCAACCGACUUUUAUAAGA |
| Mouse *Sec24c* siRNA (Cat #: M-059052-00-0005) | GCAAACGUGUGGAUGCUUA |
|  | GGUAUCAUCAGUCGAGUUA |
|  | UGGCUGAUCUGUAUCGAAA |
|  | CACCGUAUGUUGUGGAUCA |
| Mouse *Sec24d* siRNA (Cat #: M-065430-00-0005) | UAGCGGAUCUGUACAAGAG |
|  | UCGUCAAGCUCAUAUGUGA |
|  | CCAGAUACAUUCAGAUAGC |
|  | UCAAUCAGACUGCCCAUAU |
| Mouse *fibronectin 1* siRNA (Cat #: M-043446-00-0005) | GUUAGAAGCUACACCAUUA |
|  | CAACAUUGAUCGCCCUAAA |
|  | GAGAACAAACACUAACGUA |
|  | GCACCUAUCACAGGGUAUA |
| Human *SEC24A* siRNA (1) | GAGUCAGUGAGCCAAGGAUUU |
| Human *SEC24B* siRNA (Cat #: M-008299-01-0005)  Human *SEC24C* siRNA (Cat #: M-008467-01-0005)  Human *SEC24D* 5’-UTR siRNA | AUCCUUGGCUCACUGACUCUU  GGGAAAGGCUGUGACAAUA  GACCAGAAGUUCAGAAUUC  CAGGGUGCAUCUAUUAUUA  CCAGAUUCAUUCGGUGUA  GCAAACGUGUGGAUGCUUA  CAGGGAAGCUCUUUCUAUU  UGGCUGAUCUAUAUCGAAA  CUGUAUAUGAUUCGGUAUU  CGAGCGUGAGUCCAGGCUA  CGGGAGCGGGAACAGACUU |

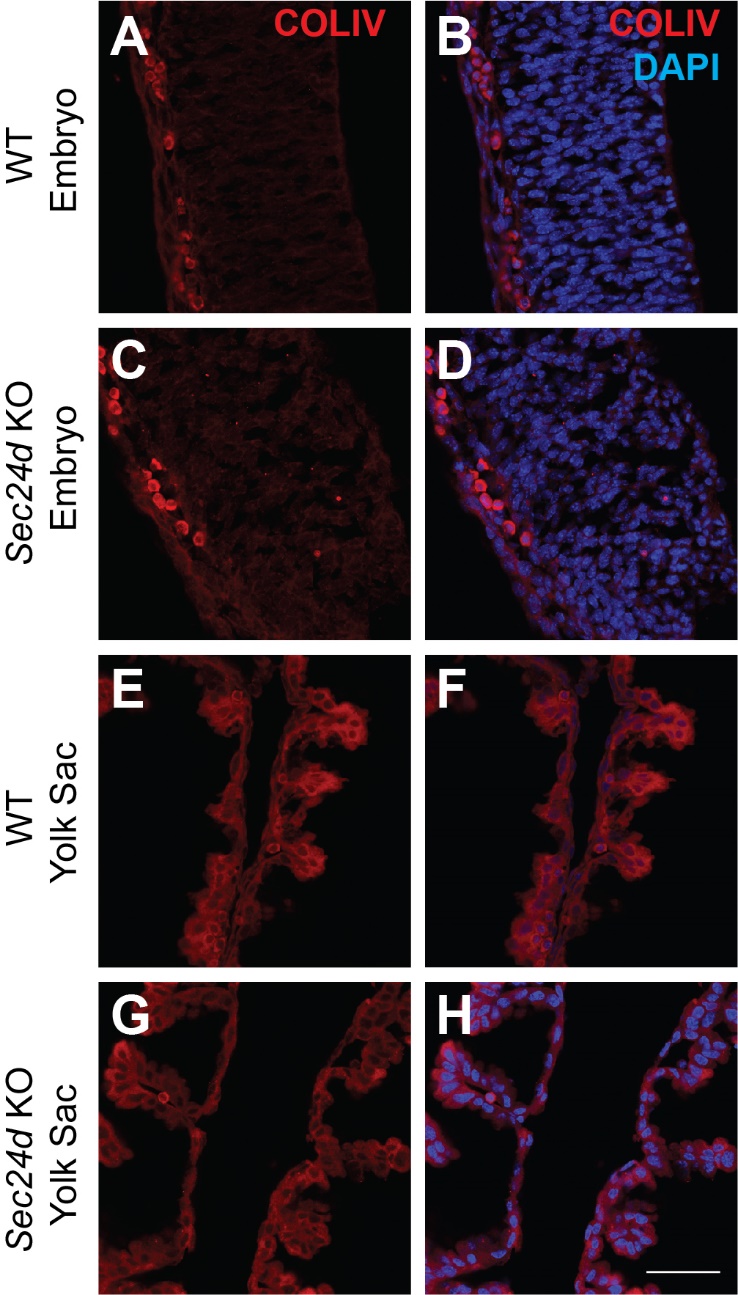

Figure S1.	Distribution of type IV collagen (COLIV). (A-H) Anti-COLIV immunoreactivity in WT (A-B, E-F) and *Sec24d* KO mutant (C-D, G-H) head skin (A-D) and yolk sac (E-H). Embryo skin and yolk sac displayed similar immunoreactivity patterns in both genotypes.

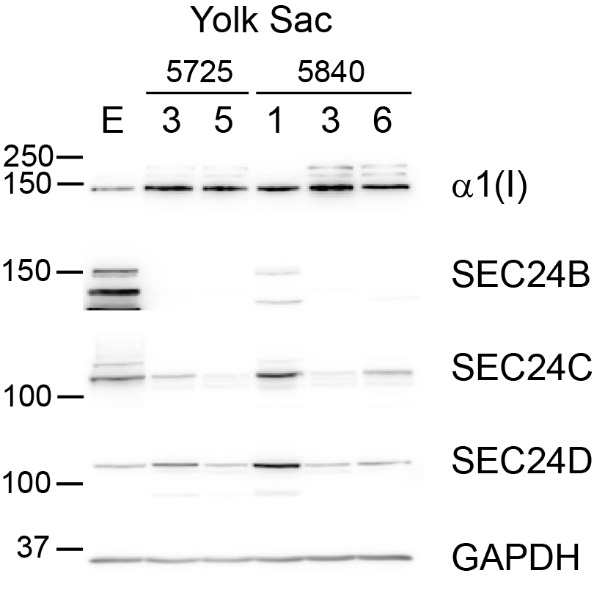

Figure S2.	Variations in the levels of SEC24s in the yolk sacs of wild-type mice. α1(I), SEC24B, SEC24C, and SEC24D were probed with immunoblotting using the yolk sac extracts of wild-type animals from litters 5725 and 5840. Embryo extracts (E) were used as reference. GAPDH was used as a loading control. While SEC24B levels are consistently low in all animals, SEC24C levels were significantly low only in 5725-5 and 5840-3 animals.

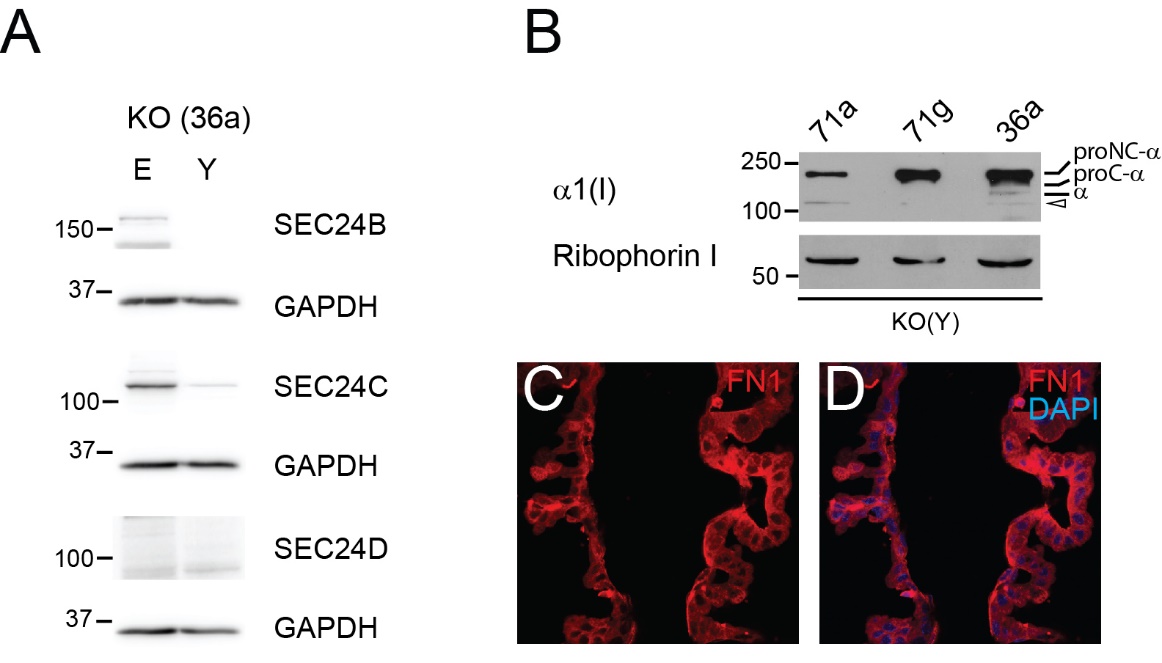

Figure S3.	Low levels of SEC24C can be observed in *Sec24d* mutant yolk sac. (A) SEC24B, SEC24C, and SEC24D were probed with immunoblotting using the extracts of a wild-type (WT) and a *Sec24d* KO mutant (36a). E, embryo; Y, yolk sac. GAPDH was used as a loading control. (B) α1(I) was probed in the yolk sac extracts of *Sec24d* KO mice (71a, 71g and 36a). Open arrowhead, abnormally processed collagen. Ribophorin I was used as a loading control. (C and D) Anti-FN1 immunoreactivity in the yolk sac of a *Sec24d* KO mutant. FN1 immunoreactivity was observed in the cells, indicating a secretion defect.

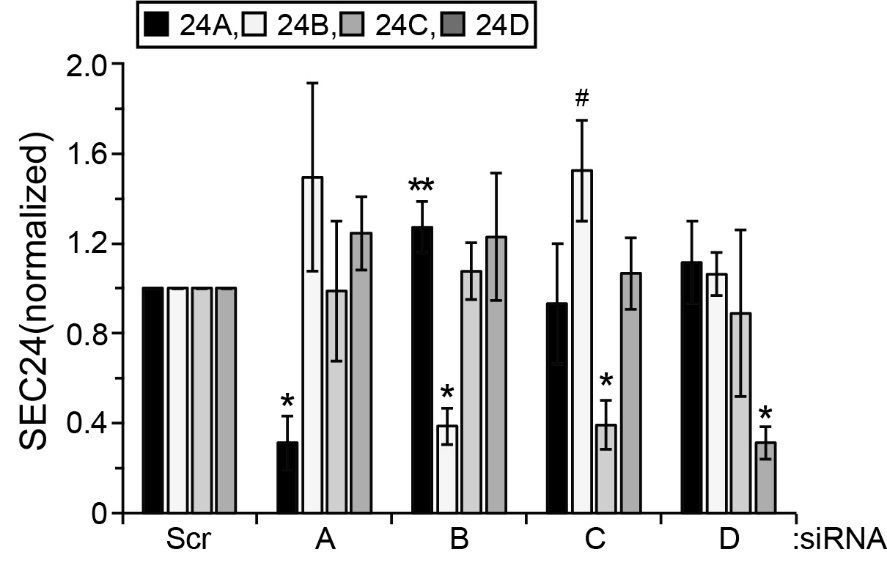

Figure S4.	Wild-type MEFs were transfected with indicated siRNAs. Statistical significance was determined with Student’s t-test. Student’s t-test was performed between a Scr control and a related SEC24 siRNA treatment (the same shade). Error bars represent standard deviations. Scr, scrambled siRNA; A, *Sec24a* siRNA; B, *Sec24b* siRNA; C, *Sec24c* siRNA. Levels of *Sec24a* mRNAs were measured with RT-qPCR and were normalized to those of *Gapdh* mRNAs. To quantify SEC24B, SEC24C and SEC24D, chemiluminescence signals from immunoblots were scanned with an Azure image scanner 600 and quantified with an AzureSpot software. The levels of SEC24 were normalized with those of α-tubulin. Then, SEC24 levels were normalized to SEC24 levels in Scr controls. When *Sec24b* expression was reduced, *Sec24a* expression was increased and when *Sec24c* expression was reduced, *Sec24b* expression was increased compared to Scr control, probably reflecting a compensatory mechanism. P*<0.0001, P**<0.05, P^#^<0.005, n=4.

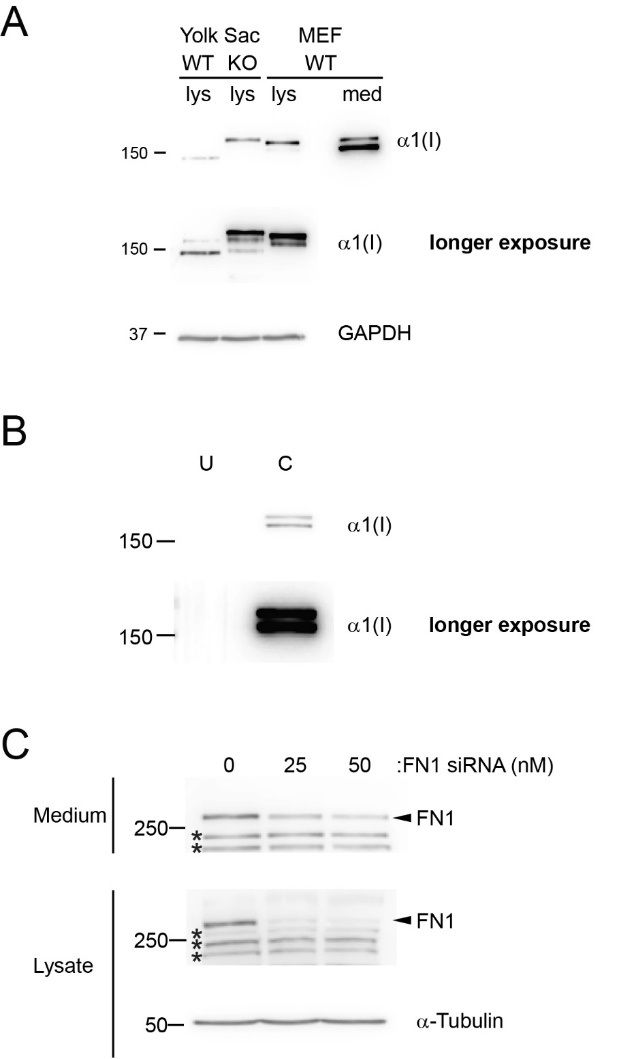

Figure S5.	(A) Collagen, α1(I), from yolk sacs of wild-type (WT) and *Sec24d* KO mice was compared with that from wild-type MEFs. lys, lysates; med, medium. Collagen species from MEFs migrated faster than those from mouse samples. This likely reflects differences in post-translational modifications (e.g., glycosylation and hydroxylation). Mature collagen was undetectable in cultured MEFs. (B) Unconditioned (U) medium (DMEM + 10% FBS) and conditioned (C) medium after 24h culture were probed for α1(I). α1(I) was not observed from the unconditioned culture medium. (C) Fibronectin 1 knockdown. Wild-type MEFs were transfected with fibronectin 1 (FN1)-specific siRNAs at day 0. The culture medium was replaced with fresh medium at day 1. Cells and conditioned media were harvested at day 2. Cell lysates and media were analyzed by immunoblotting. Asterisks, nonspecific proteins.

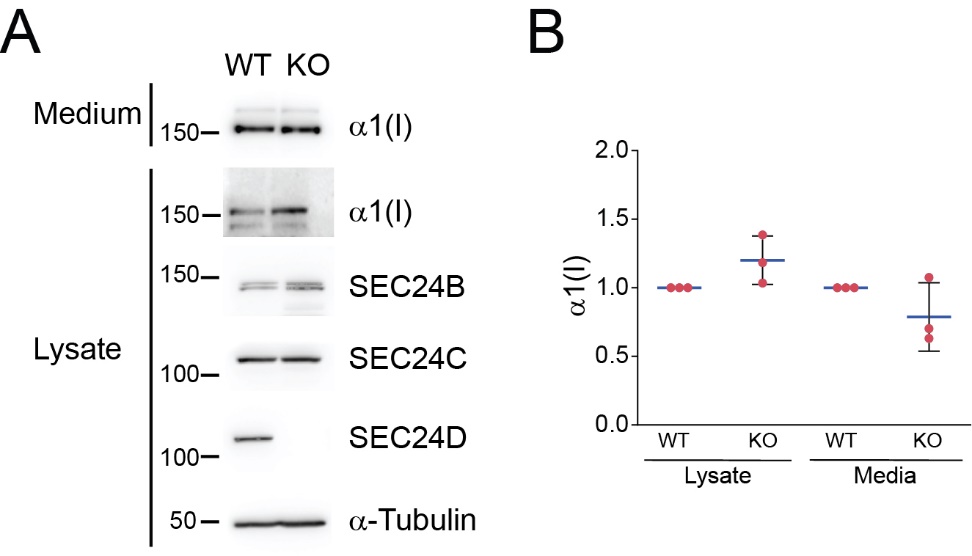

Figure S6.	Secretion of collagen, α1(I), in wild-type and *Sec24d* KO MEFs. Confluent cells were incubated with fresh medium and incubated for 24h. Cells and conditioned media were collected and processed for immunoblotting. (A) A representative immunoblot. (B) A summary of three independent experiments. Levels of α1(I) were normalized to those of α-tubulin. Statistical significance was determined with Student’s t-test. Error bars represent standard deviations (n=3). WT, wild-type; KO, *Sec24d* knockout.

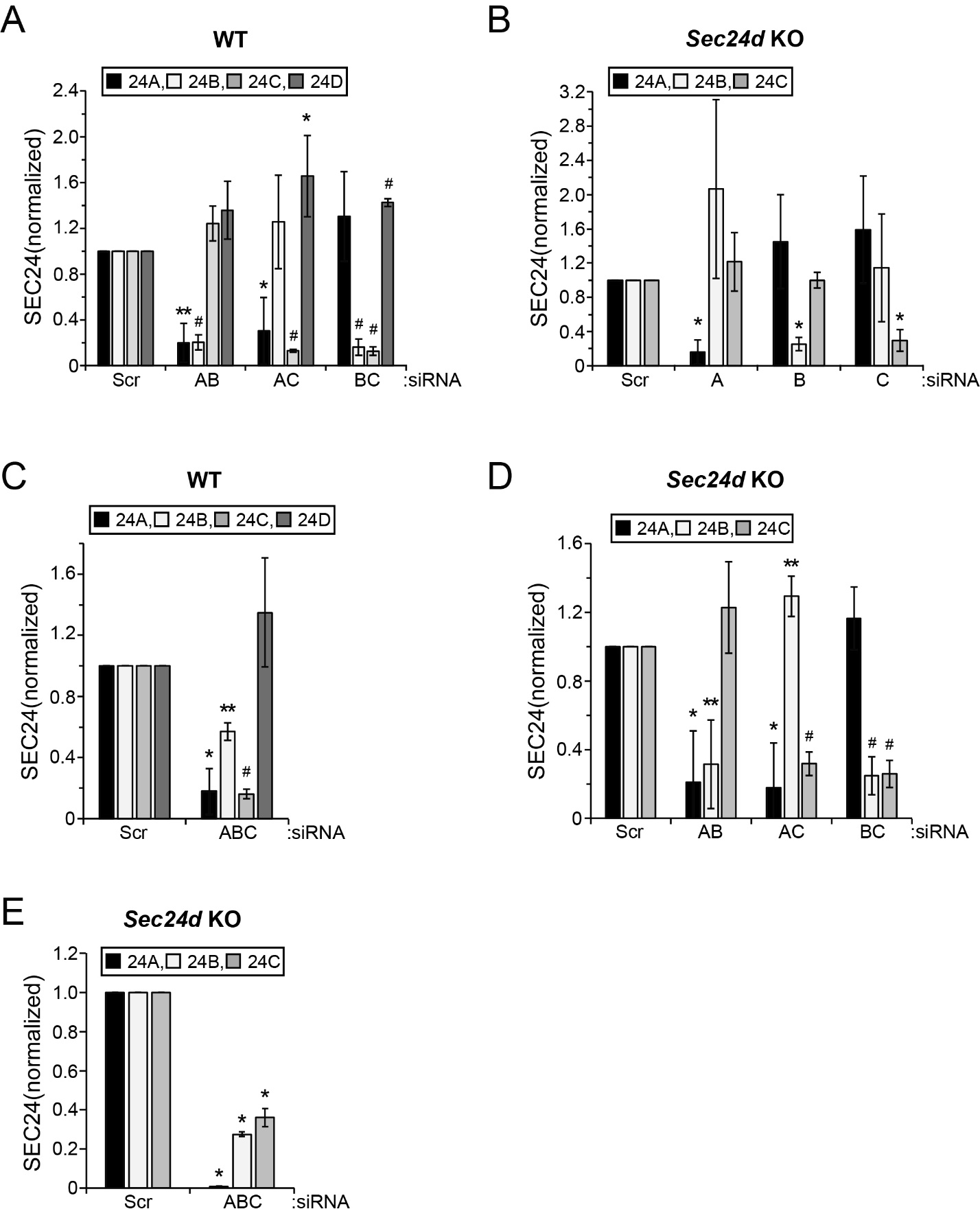

Figure S7.	Wild-type (A and C) and *Sec24d* KO (B, D and E) MEFs were transfected with indicated siRNAs. Statistical significance was determined with Student’s t-test. Student’s t-test was performed between a Scr control and a related SEC24 siRNA treatment (the same shade). Error bars represent standard deviations. Scr, scrambled siRNA; A, *Sec24a* siRNA; B, *Sec24b* siRNA; C, *Sec24c* siRNA. Levels of *Sec24a* mRNAs were measured with RT-qPCR and were normalized to those of *Gapdh* mRNAs. To quantify SEC24B, SEC24C and SEC24D, chemiluminescence signals from immunoblots were scanned with an Azure image scanner 600 and quantified with an AzureSpot software. The levels of SEC24 were normalized with those of α-tubulin. Then, SEC24 levels were normalized to SEC24 levels in Scr controls. (A) P*<0.05, P**<0.005, P^#^<0.0001, n=4. (B) P*<0.0001, n=4. (C) P*<0.001, P**<0.0005, P^#^<0.0001, n=4. (D) P*<0.01, P**<0.005, P^#^<0.0001, n=4. (E) P*<0.0001, n=3.

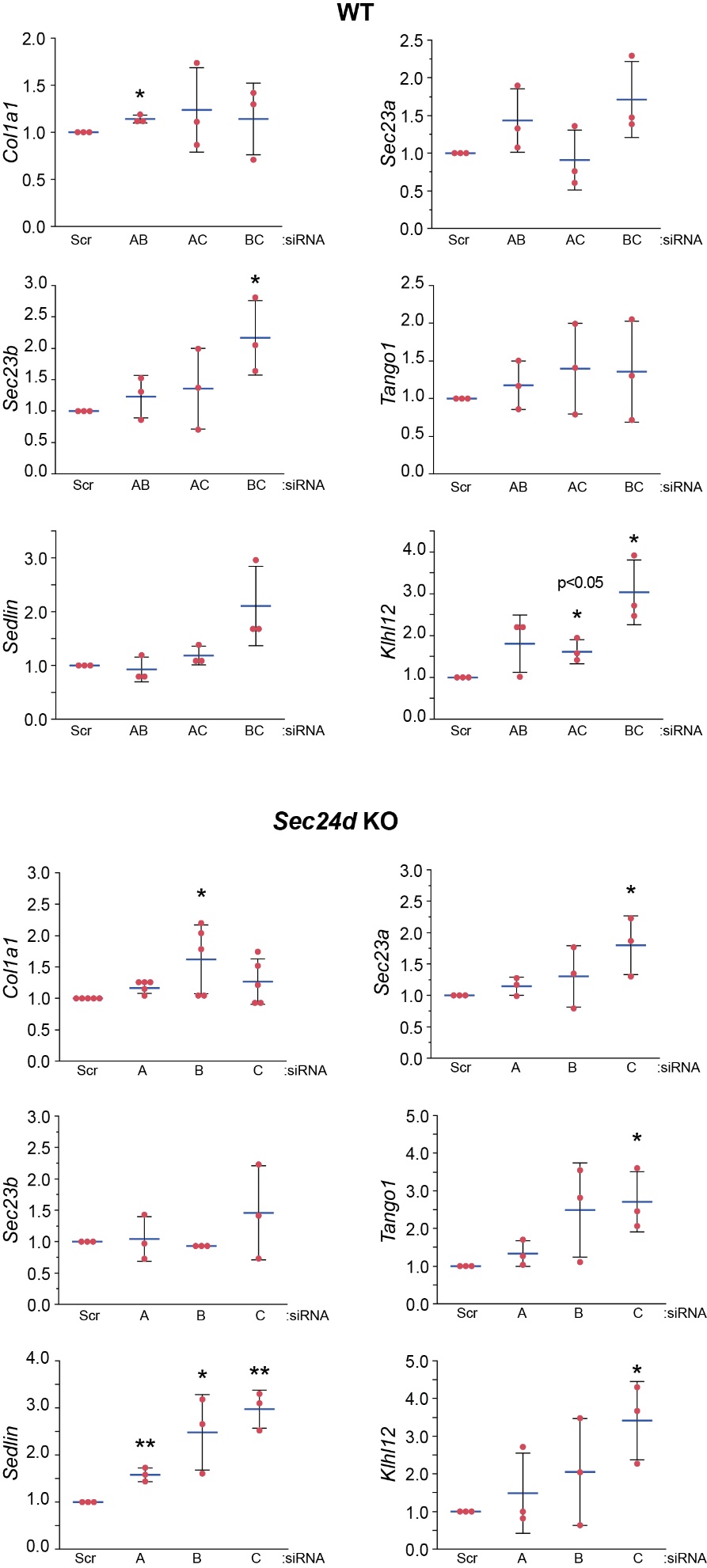

Figure S8.	Changes in the expression of collagen and collagen-related genes in wild-type (WT) and *Sec24d* KO MEFs. Scr, scrambled siRNA; A, *Sec24a* siRNA; B, *Sec24b* siRNA; C, *Sec24c* siRNA. Levels of indicated transcripts were measured with RT-qPCR and were normalized to those of *Gapdh* mRNAs. Student’s t-test: P*<0.05, P**<0.005, n was indicated in dots. Error bars represent standard deviations and middle lines represent averages.

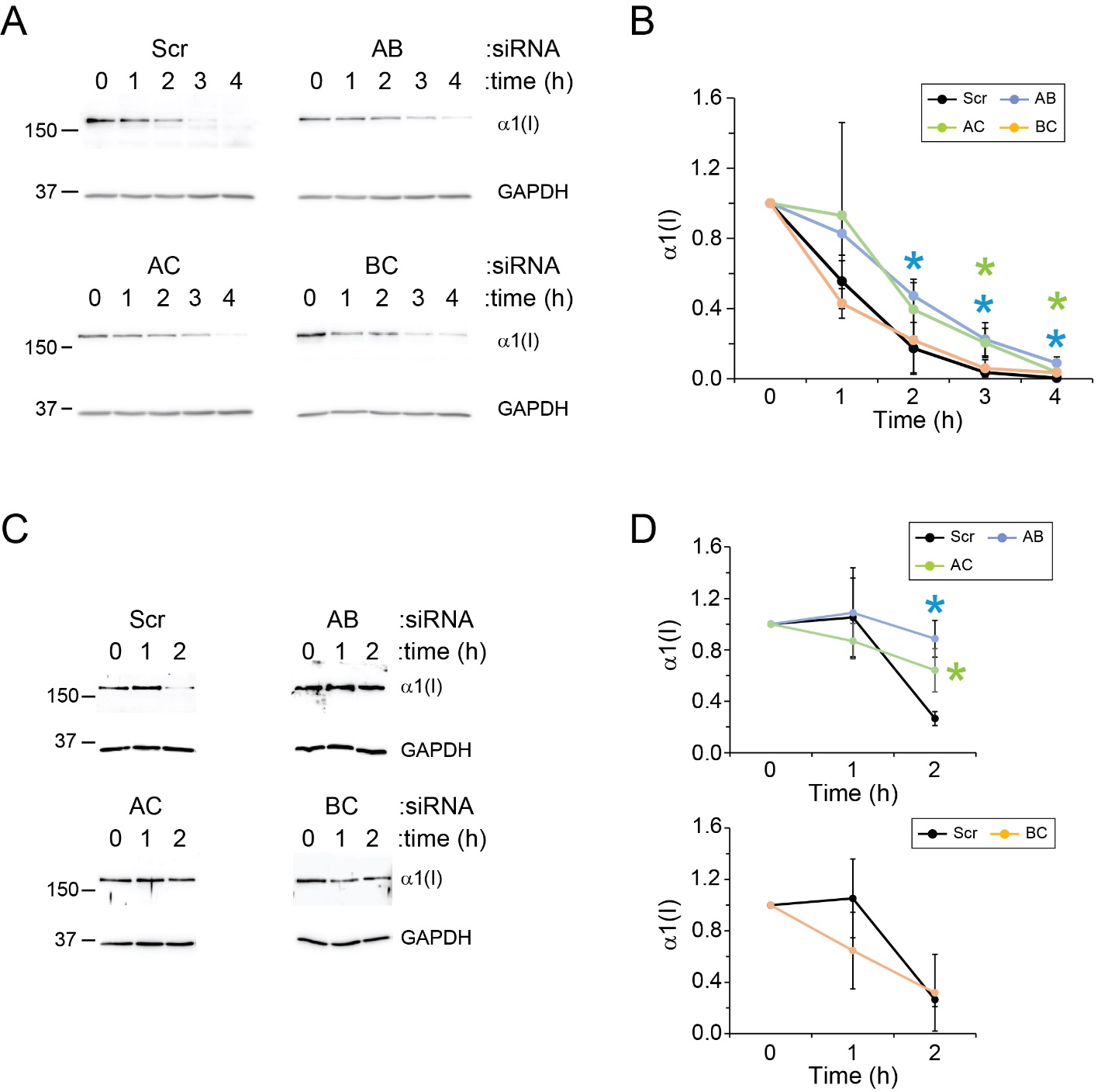

Figure S9.	Delayed clearance of cellular collagen. Wild-type MEFs were transfected with indicated SEC24 siRNAs. In the following day, the cells were treated with cycloheximide (100 μg/ml) (A and B) or cycloheximie and bafilomycin A1 (100 μg/ml and 50 nM, respectively) (C and D). The cells were sampled at indicated times. Scr, scrambled siRNA; A, *Sec24a* siRNA; B, *Sec24b* siRNA; C, *Sec24c* siRNA. Levels of cellular collagen were normalized to those of GAPDH. (B) Student’s t-test compared to the scrambled (Scr) siRNA condition: AB, P*<0.05, n=3; AC, P*<0.05, n=3. Error bars represent standard deviations. (D) Student’s t-test compared to the scrambled (Scr) siRNA condition: AB, P*<0.005, n=3; AC, P*<0.05, n=3. Error bars represent standard deviations.

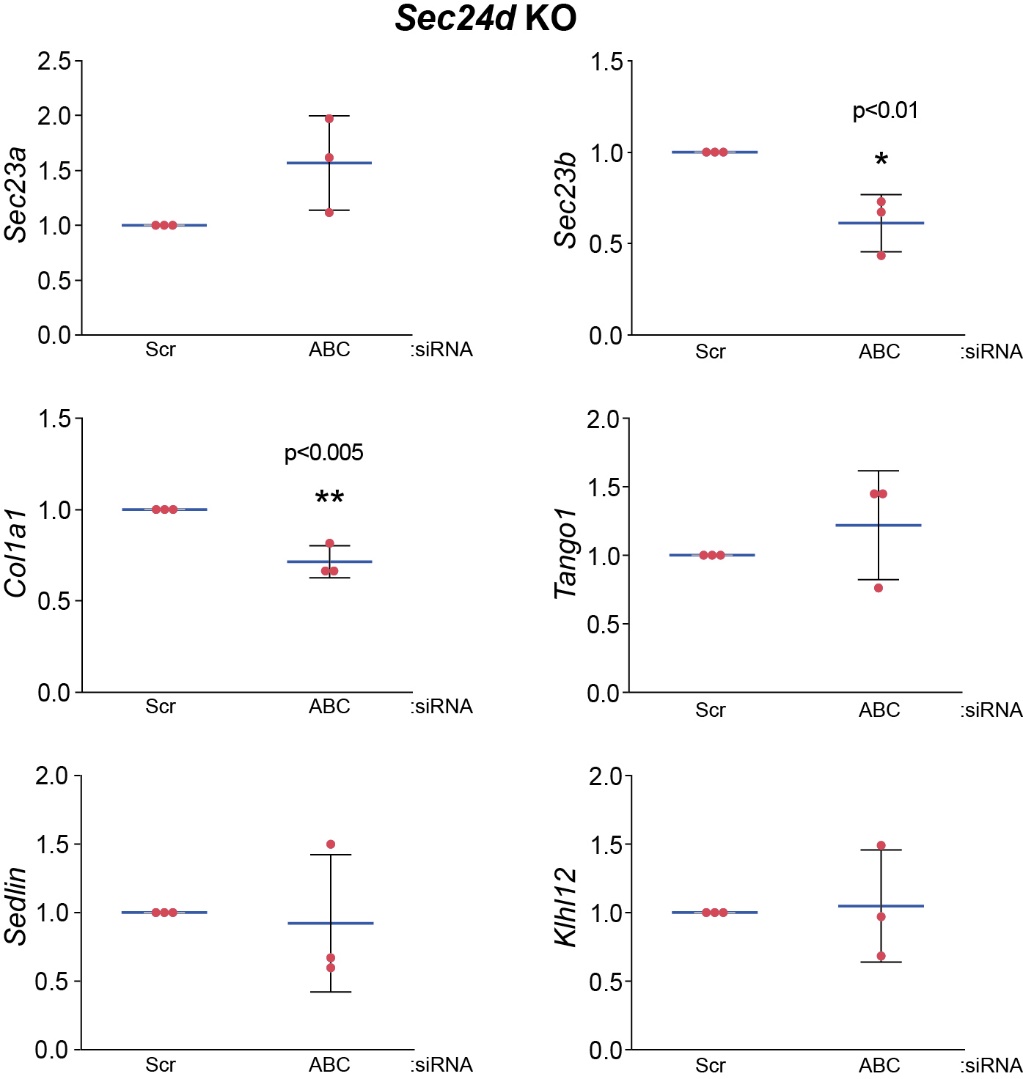

Figure S10.	Changes in the expression of collagen and collagen-related genes in *Sec24d* KO MEFs. Scr, scrambled siRNA; A, *Sec24a* siRNA; B, *Sec24b* siRNA; C, *Sec24c* siRNA. Levels of indicated transcripts were measured with RT-qPCR and were normalized to those of *Gapdh* mRNAs. Student’s t-test: P*<0.01, P**<0.005, n=3. Error bars represent standard deviations and middle lines represent averages.

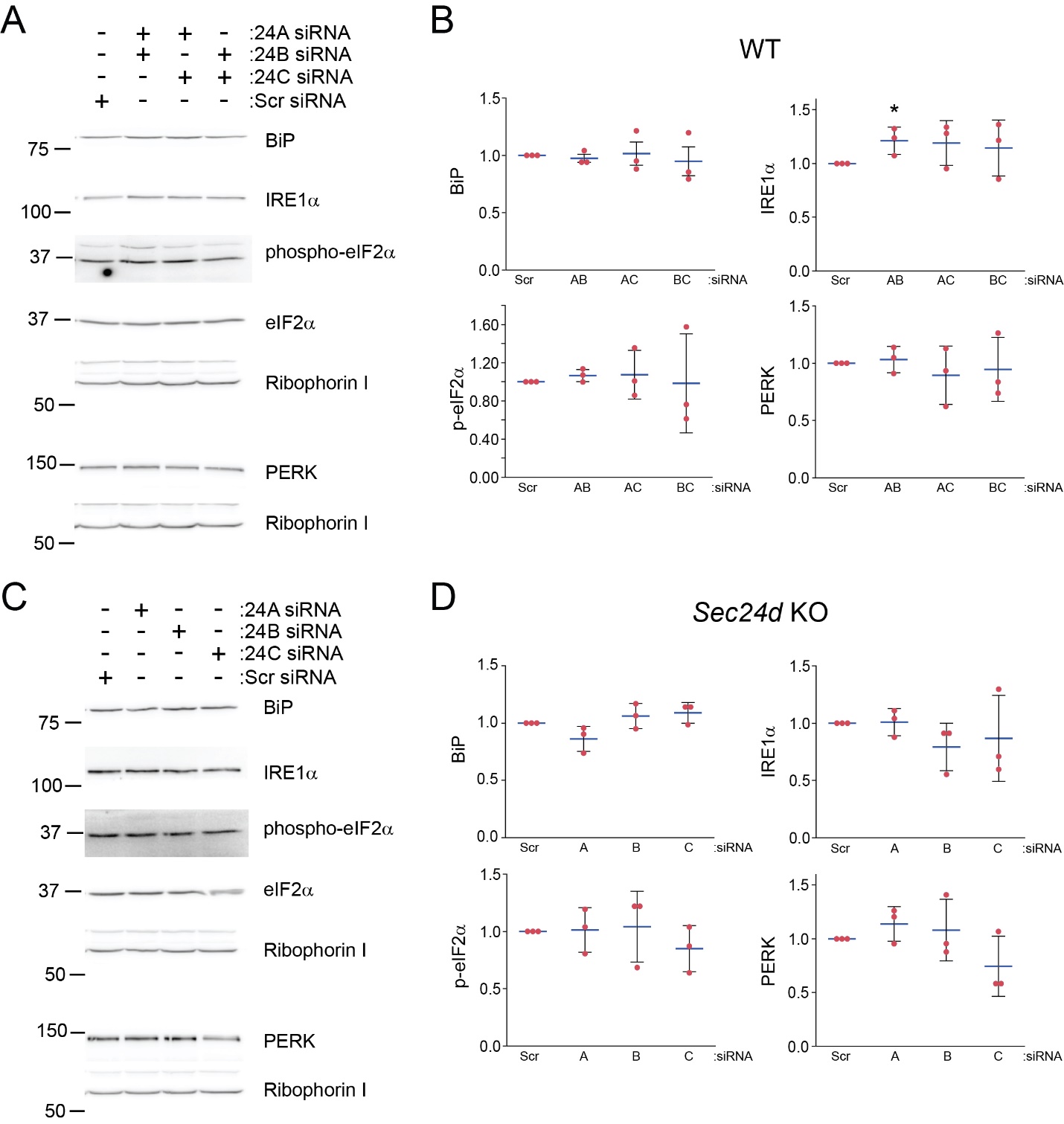

Figure S11.	Minimal dysregulation of the unfolded protein response (UPR) of the ER. Wild-type (A and B) and *Sec24d* KO (C and D) MEFs were transfected with indicated siRNAs. Scr, scrambled siRNA; A, *Sec24a* siRNA; B, *Sec24b* siRNA; C, *Sec24c* siRNA. Levels of indicated proteins were normalized to those of ribohporin I or eIF2α (for p-eIF2α). Student’s t-test: P*<0.05, n=3. Error bars represent standard deviations and middle lines represent averages.

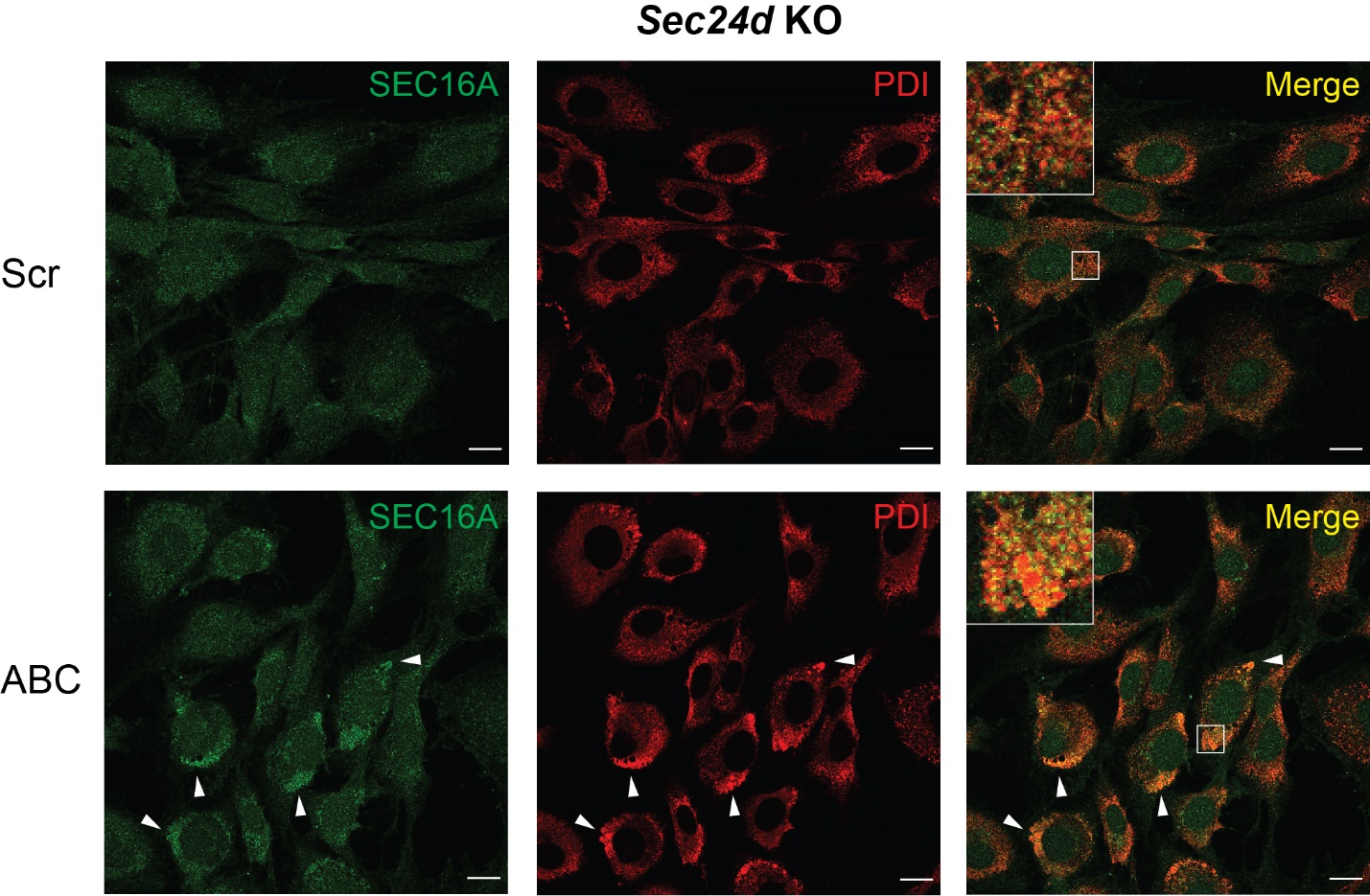

Figure S12.	Immunofluorescent micrographs. *Sec24d* KO MEFs were transfected with indicated siRNAs and immunostained ­for SEC16A and PDI. Scale bar, 20 μm. Inset, an enlarged image of the indicated region with a box. Arrowheads, a region of the ER that shows strong SEC16A signals.

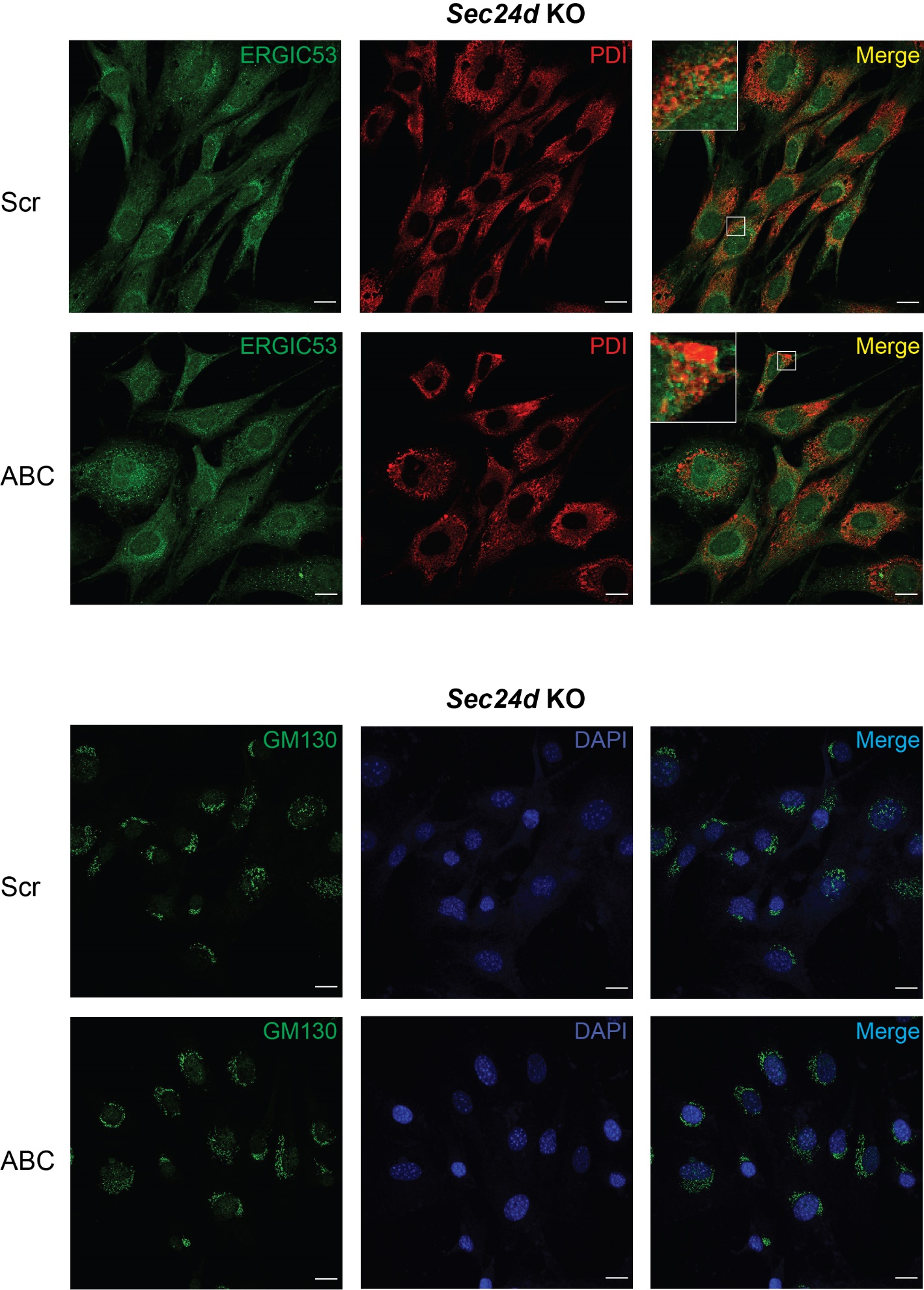

Figure S13.	Immunofluorescent micrographs. *Sec24d* KO MEFs were transfected with indicated siRNAs and immunostained for ERGIC53, GM130, and PDI. Scale bar, 20 μm.

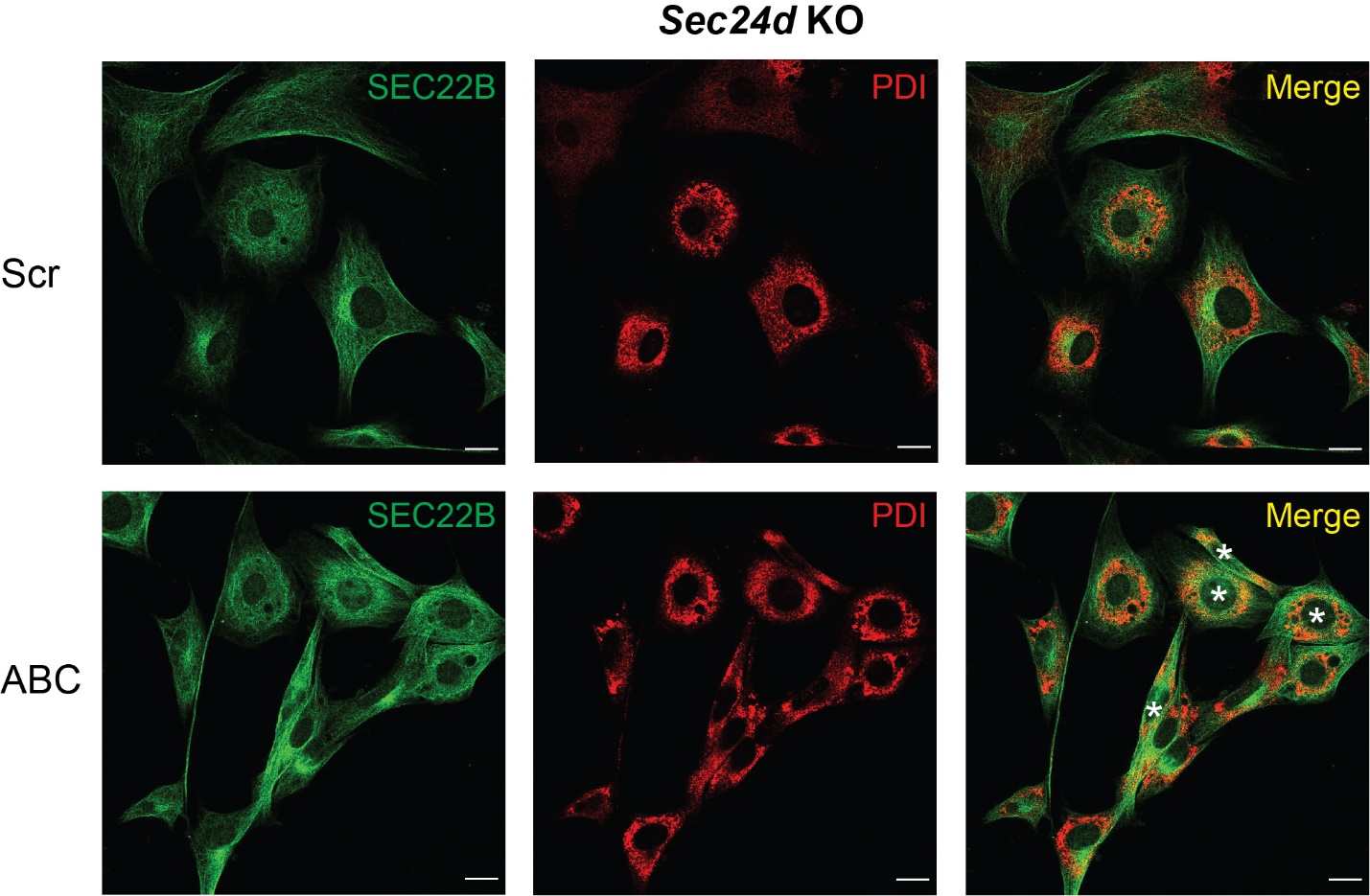

Figure S14.	Immunofluorescent micrographs. *Sec24d* KO MEFs were transfected with indicated siRNAs and immunostained for SEC22B and PDI. Cells showing extensive overlap between SEC22B and PDI were marked with asterisks. Scale bar, 20 μm.

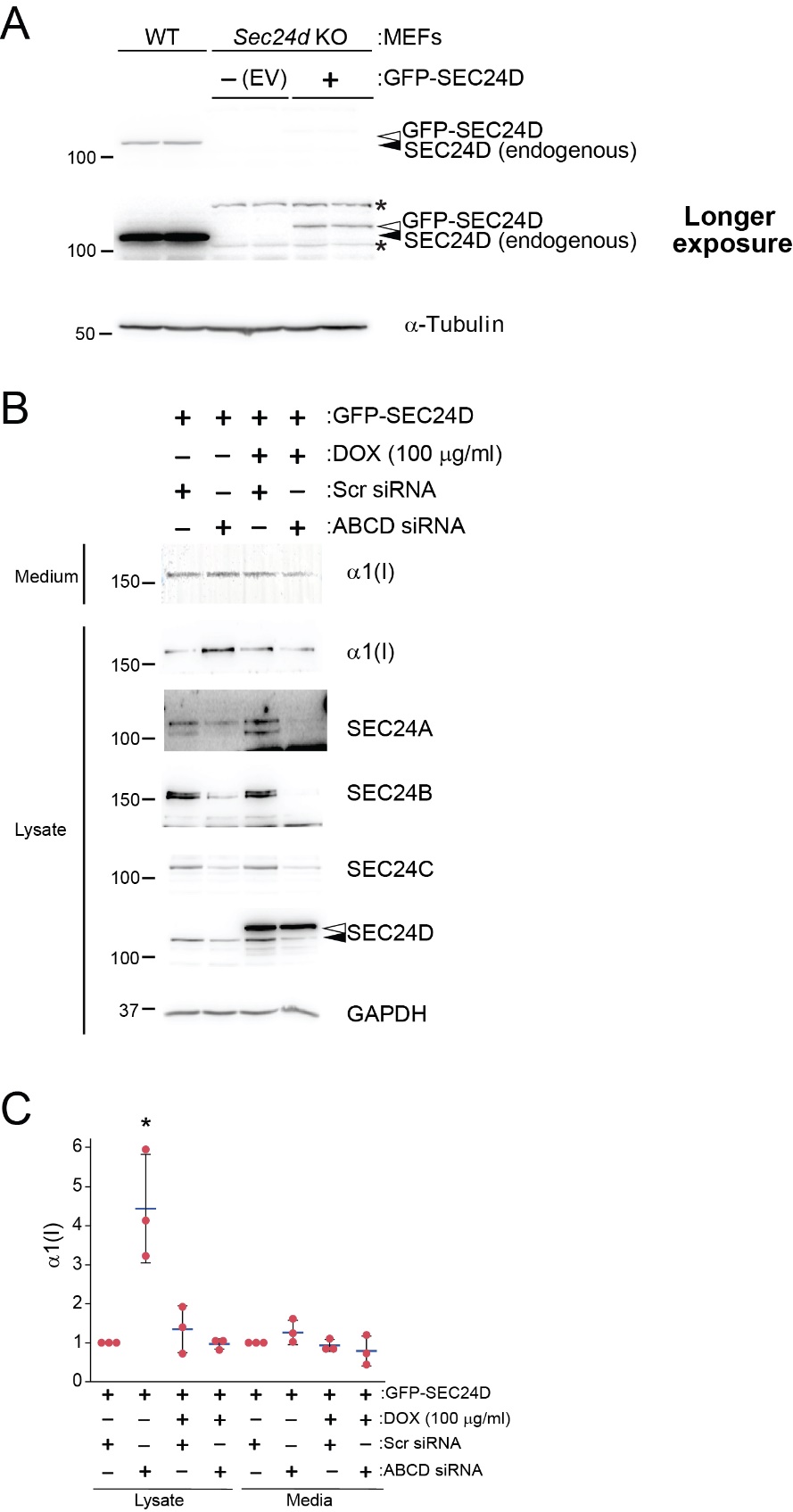

Figure S15.	A recombinant human *SEC24D* rescued the cellular collagen accumulation defect in U2OS cells. (A) *Sec24d* KO MEFs were infected with lentiviruses harboring pLVX-TRE3G-GFP-SEC24D or empty vector (EV). Infected cells were treated with puromycin to enrich cells containing the GFP-SEC24D construct. The levels of the recombinant *SEC24D* (open arrowhead) were much lower than those of endogenous SEC24D (closed arrowhead) in MEFs. (B and C) U2OS cells were co-infected with lentiviruses harboring pLVX-Tet3G and pLVX-TRE3G-GFP-SEC24D. Infected cells were selected with G418 and puromycin to generate a stable cell line containing the Tet-inducible *GFP-SEC24D* construct. The recombinant *SEC24D* (open arrowhead) were overexpressed, compared to endogenous SEC24D (closed arrowhead) in a doxycycline-dependent manner. The stable cells were transfected with indicated siRNAs targeting human SEC24s. Scr, scrambled siRNA; A, *SEC24A* siRNA; B, *SEC24B* siRNA; C, *SEC24C* siRNA; D, *SEC24D* siRNA. *SEC24D* siRNAs target 5’ UTR of *SEC24D* mRNAs (Table S2). Levels of indicated proteins were normalized to those of GAPDH. Student’s t-test: P*<0.05, n=3. Error bars represent standard deviations and middle lines represent averages.
